# Genetic and functional diversity of allorecognition receptors in the urochordate, *Botryllus schlosseri*

**DOI:** 10.1101/2024.10.16.618699

**Authors:** Henry Rodriguez-Valbuena, Jorge Salcedo, Olivier De Their, Jean Francois Flot, Stefano Tiozzo, Anthony W De Tomaso

**Author notes:** **Corresponding author:** Anthony De Tomaso.

## Abstract

Allorecognition in *Botryllus schlosseri* is controlled by a highly polymorphic locus (the *fuhc*), and functionally similar to missing-self recognition utilized by Natural Killer cells-compatibility is determined by sharing a self-allele, and integration of activating and inhibitory signals determines outcome. We had found these signals were generated by two *fuhc*-encoded receptors, called *fester* and *uncle fester*. Here we show that *fester* genes are members of an extended family consisting of >37 loci, and co-expressed with an even more diverse gene family-the *fester co-receptors* (*FcoR*). The *FcoRs* are membrane proteins related to *fester*, but include conserved tyrosine motifs, including ITIMs and hemITAMs. Both genes are encoded in highly polymorphic haplotypes on multiple chromosomes, revealing an unparalleled level of diversity of innate receptors. Our results also suggest that ITAM/ITIM signal integration is a deeply conserved mechanism that has allowed convergent evolution of innate and adaptive cell-based recognition systems.

## Introduction

Highly polymorphic allorecognition systems are found in species throughout the metazoa, and studies on allogeneic transplantation led to the discovery of both adaptive and innate cell-based recognition systems in vertebrates^1,2^. Discriminatory ability in adaptive immunity is based on somatic recombination and thymic selection, identifying TCRs with structures that can discriminate between even a single residue on a pMHC complex ^3,4^. Innate allorecognition is mediated by NK cells, which tally binding from a repertoire of stochastically expressed activating and inhibitory receptors to determine the health of a target cell^5^. Inhibitory receptors bind to polymorphic epitopes on MHC Class I allotypes, allowing NK cells to determine their presence or absence, a strategy called missing-self recognition^2^

Both innate and adaptive receptors utilize the same signaling pathways-the TCR and NK activating receptors use the ITAM pathway, while inhibitory receptors in both cells use the ITIM pathway, and discrimination is due to an interaction between kinase and phosphatase activity ^6–8^. Finally, during development both cells go through a cell autonomous education process, which quantifies binding potential and sets an activation threshold that allows cells to be self-tolerant and functional^4,9,10^.

Interestingly, T-cell recognition appears as a functioning unit in jawed vertebrates-but no homologs of the MHC, TCR, RAG, or evidence of thymic selection have ever been identified in jawless fish, non-vertebrate chordates, or invertebrates ^11,12^. NK receptors are evolving even faster: rodents use the C-type lectin *Ly49* genes, while the primate analogs are the Killer cell immunoglobulin-like receptors (*KIRs*)^13^- and neither are found outside of the mammals^14^. This plasticity is not restricted to vertebrates-histocompatibility has been characterized in three non-vertebrate clades, and in each, candidate genes also emerge abruptly, with no homologs found outside of closely related species^15–17^. However, both vertebrate and candidate non-vertebrate allorecognition receptors are part of multigene families encoded in polymorphic haplotypes^14,15,18,19^, and activating and inhibitory members encoding ITAM and ITIM signaling motifs are found in both the cnidarian, *Hydractinia*^19^ and *Botryllus* (this study). The recurring evolution of both vertebrate and non-vertebrate recognition systems that all converge onto equivalent signaling pathways suggest that the cellular mechanisms that process information from cell surface binding events are conserved, allowing receptors to evolve freely^20^.

*Botryllus schlosseri* is a urochordate, the sister group to the vertebrates^21^, and provides a novel model to investigate mechanisms underlying polymorphic discrimination and the evolution of vertebrate immunity. Botryllus undergoes a natural transplantation reaction when two individuals grow into proximity. Terminal projections of the vasculature, called ampullae come into contact. If two individuals are compatible, the ampullae will *fuse*, forming a parabiosis. If they are incompatible, the vessels will *reject*, an inflammatory reaction which prevents vascular fusion (Extended Data Fig. 1). Fusion or rejection is controlled by a single, highly polymorphic locus called the *fuhc* (**fu**sion/**h**isto**c**ompatibility) and discrimination occurs by missing-self recognition-individuals are compatible if they share one or both *fuhc* alleles, but are incompatible if no alleles are shared ^11,22^.

Botryllus populations have 200-500 fuhc alleles, thus the effector system can pinpoint a self-allele from hundreds of competing nonself alleles^23,24^, but the mechanisms that allow an innate recognition system to carry out this level of discrimination are not understood. We had previously characterized two histocompatibility receptors encoded in the fuhc locus: *fester* is highly polymorphic and required for fusion, while *uncle fester* is monomorphic and is required for rejection^25,26^. Similar to NK cell recognition, the outcome is due to an interaction between these two independent pathways, however, neither gene encodes known signaling domains, and it was unclear if other receptors contributed to specificity.

Here we find that rather than being encoded in only two loci, the *fester* genes are part of a large, multigene family, and co-expressed with members of another multigene family-the *fester coreceptors (FcoR)*. FcoR are type I transmembrane proteins that encode ITIM and hemITAM motifs. These results suggest that two families of immune receptors participate in the *Botryllus* histocompatibility response, and that polymorphic discrimination requires an interaction between ITAM and ITIM signaling.

## Results

### *fester* and *uncle fester* are members of a large gene family

Fester and uncle fester share a similar domain structure (Fig. 1a,c), but amino acid similarity is only found in the COOH half of the protein, encoding the TM and intracellular domains. We identified low homology structural homologs of fester in the fuhc locus of a related ascidian^27^, and used those to search in 26 transcriptome and four genome databases of Botryllus (Extended Data Table 1), and identified 45 unique sequences (Fig. 1a, b Extended Data Fig. 2a). On average, these sequences shared 20% identity at the amino acid level (Extended Data Fig. 2a, b), which was concentrated in the COOH tail half of the protein.

**Fig. 1.**
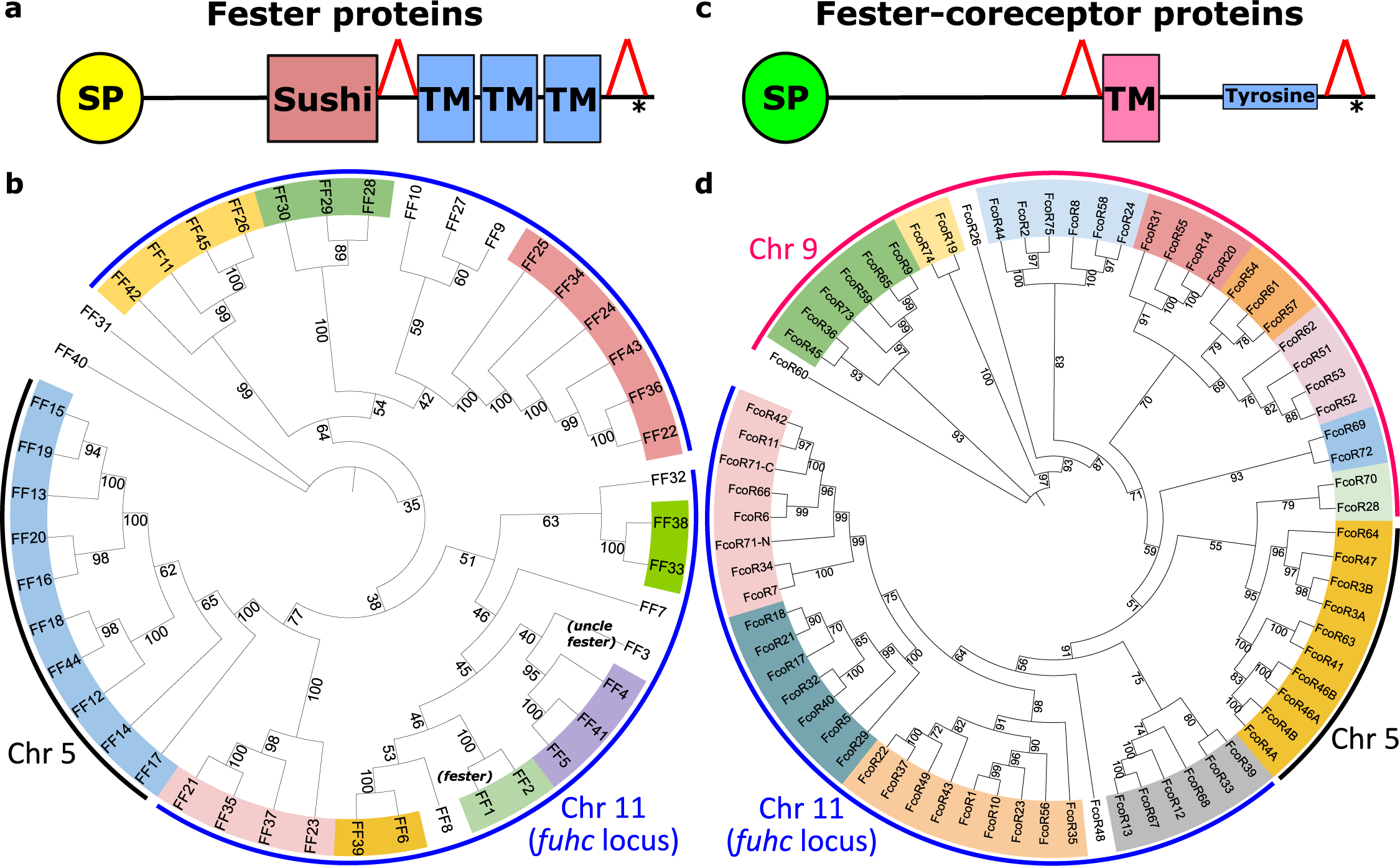
Phylogenetic analysis of full-length Fester and FcoR proteins. **a,c** Illustration of domain architectures and **b,d** phylogenetic trees of the full-length fester (n=45) and FcoR proteins (n=69) performed using Maximum likelihood method. Bootstrap supports are indicated at the base of the branches. Well-supported clades are highlighted in different colors. *FF1* (*fester*) and *FF3* (*uncle fester*) genes are indicated. Putative chromosomal localizations are indicated (Fig. 4). Red triangles in a,c illustrate regions that are alternatively spliced.

To estimate which of these sequences were alleles vs loci, we used the range of protein identity values from a population genetic study of *fester* as a benchmark (Extended Data Fig. 2b) ^28^. If sequences shared >82% amino acid homology and >90% nucleotide homology over the entire gene, they were classified as alleles, predicting 37 new loci. We are calling these new genes the fester family (FF), and *fester* and *uncle fester* have been renamed FF1 and FF3, respectively. Phylogenetic analysis (Fig. 1b) sorted these loci into multiple clades. Each individual expressed a nearly unique repertoire of fester genes, with an average of 10 fester genes or alleles per individual, with a range of 7-13 (Extended Data Fig. 3, Table 3), suggesting that FF loci are encoded in complex haplotypes with presence/absence polymorphisms.

### Genetic properties of the fester family members

The genes encoding the FF1 and FF3 are composed of eleven and nine exons, respectively (Extended Data Fig. 4a). Each gene shares a common overall structure: exon one encodes a signal sequence, a Sushi domain is encoded on exon 5 (FF1) and 3 (FF3), and in both cases the Sushi domain is followed by two short exons that encode extracellular regions, followed by three exons, each encoding one transmembrane domain ^25,26^. We determined the genomic structure for nine other fester family members, and they are encoded in 9, 10, or 12 exons, with a similar exon architecture in the center of the gene as described above. (Extended Data Fig. 4a).

We had previously found that FF1 is highly polymorphic, identifying 60 allotypes in the U.S^28^. In contrast, FF3 was nearly monomorphic, as only two alleles that differed by a single amino acid were identified^26^. Of the 37 new loci, three are polymorphic (FF12, FF13 and FF28), but not to the level of FF1. These three loci each have 3-4 alleles that are ca. 90% identical. In contrast, 14 of the new loci were clearly monomorphic, as identical nucleotide sequences were identified in > 5 individuals.

However, as almost half of the unique sequences were present in only one or two individuals (Extended Data Table 3), this is likely an underestimate. Nevertheless, many of the new loci are monomorphic.

### Alternative splicing in the extracellular and intracellular regions of fester genes

One common characteristic of the known fester genes was the alternative splicing of two extracellular exons following the Sushi domain, moving that exon relative to the plasma membrane (Fig. 1a). We found that 26 of the new loci showed equivalent patterns in the extracellular region (Extended Data Fig. 4a and Table 5). Alternative splicing was also detected in the cytoplasmic tail of 12 fester genes (Extended Data Fig. 4a and Table 5). These splice variants generate new stop codons, changing the structure of the cytoplasmic tail, which may affect interactions with other proteins (described below).

### FF genes diversify by intergenic recombination

Fester family members have high similarity in their C-terminal regions, but low similarity in their extracellular regions (Fig 1a). To characterize the evolution of these regions, we analyzed the extracellular and intracellular (transmembrane and cytoplasmic) regions of the fester proteins independently (Fig. 2a,b). Analysis of the intracellular region identified five well supported clades (Fig. 2b). However, most of the ectodomains of the same fester loci split into different clades when analyzed independently (Fig. 2a), suggesting that the fester genes are diversifying by intergenic recombination, mixing and matching the ectodomains and intracellular domains. However, one clade of fester genes did not exhibit intergenic recombination (orange in Fig. 2a,b). As discussed below, this is a result of the presence of two separate clusters of FF genes, restricting the recombination of extracellular and intracellular regions to the genes encoded on the same chromosome (Fig. 4).

**Fig. 2.**
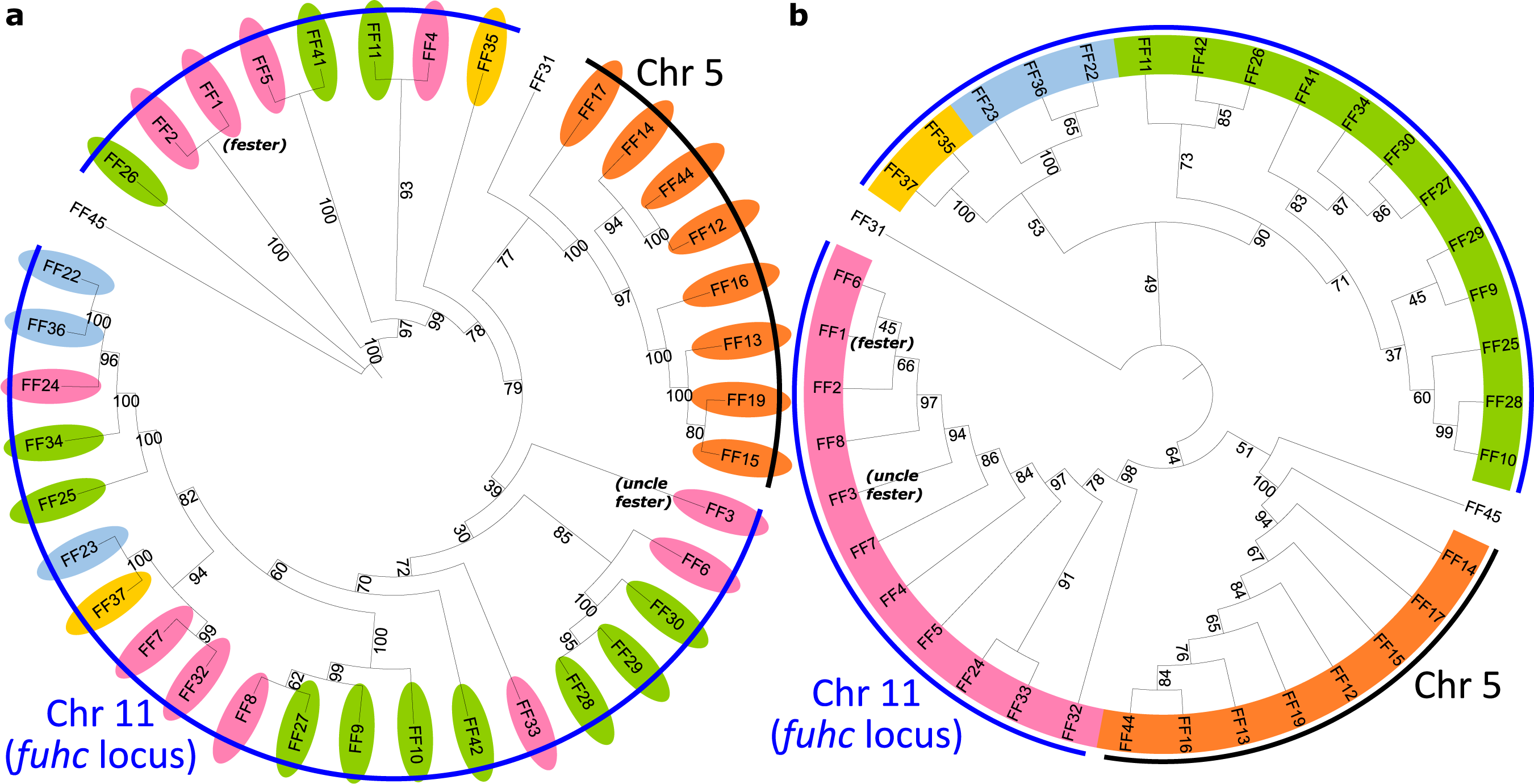
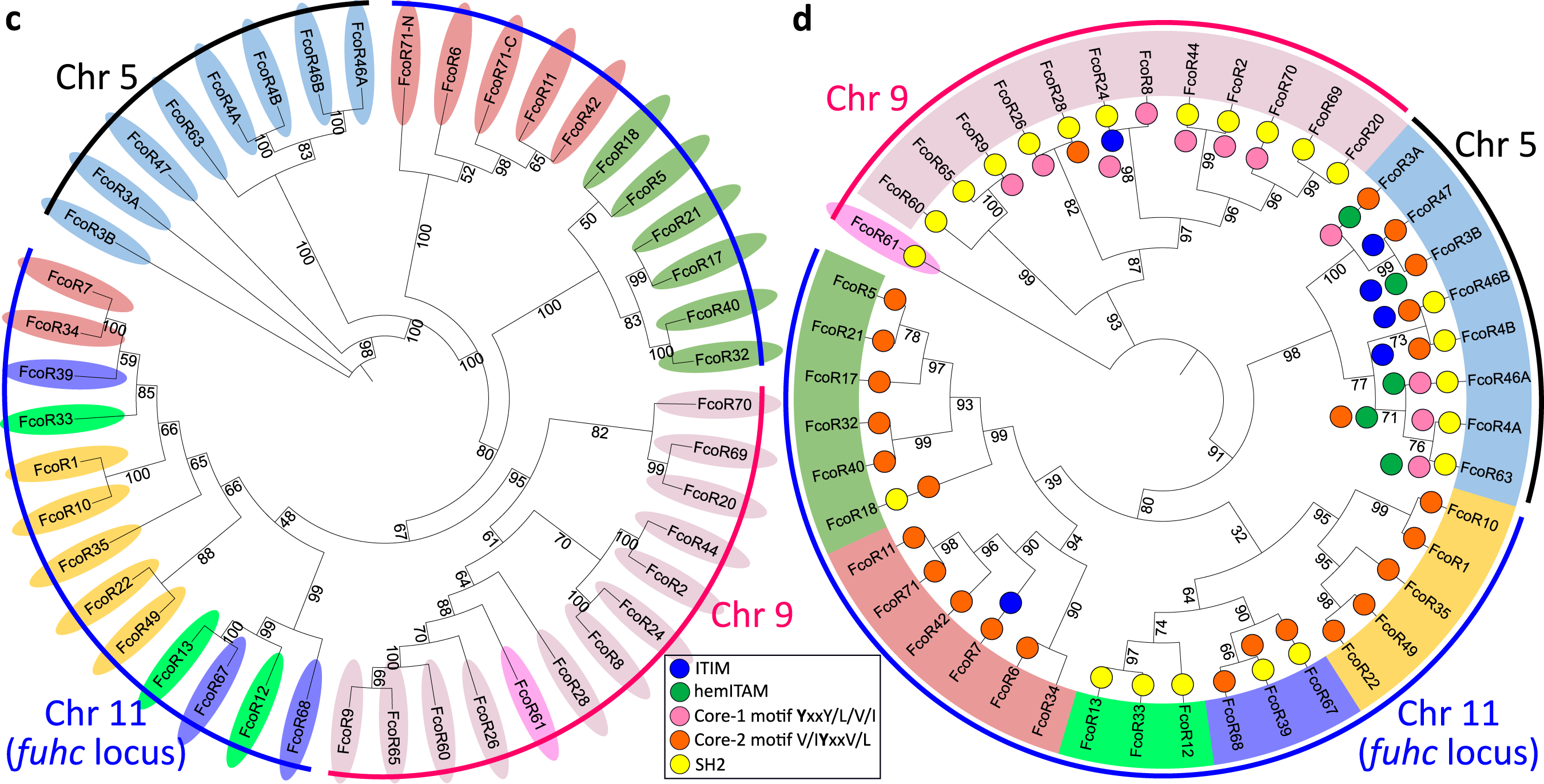
Phylogenetic trees of extracellular and intracellular regions of FF and FcoR proteins. **a,** Phylogenetic trees of the extracellular and **b,** intracellular regions of Fester proteins (n=38). **c,** Phylogenetic trees of the extracellular and **d,** intracellular regions of FcoR (n=44). Bootstrap supports are indicated at the base of the branches. Clades were highlighted based on their similarity in the intracellular regions. Tyrosine-based motifs for FcoR proteins are indicated (colored circles).

### Identification of the fester co-receptor (FcoR) gene family

Our search for new fester genes also identified another group of conserved sequences located in the fuhc locus. These genes encoded a type I protein with a single transmembrane domain (Fig. 1c; Extended Data Fig. 4b), and ranged in size from 520-610 aa. They also encoded canonical ITIMs, hemITAMs and other signaling motifs in their cytoplasmic tails. We are tentatively naming these genes the Fester co-receptors (FcoRs).

The FcoR genes are even more diverse than the FF genes, and we identified 69 unique sequences in the same databases (Extended Data Fig 2c). Nine were full length, encoding a signal peptide and stop codon, 37 were nearly fully length and had a predicted TM, intracellular tail and most had a stop codon (Extended Data Fig. 6). We also found 23 partial sequences that overlapped with the extracellular region of the full-length sequences. As described below, one of these loci (FcoR7) was highly polymorphic, and we used the same analysis as for fester to estimate that there are 53 unique FcoR loci (Extended Data Fig. 2c,d). Each individual expressed an almost unique repertoire of FcoR genes (Extended Data Fig. 8; Table 3), with an average of 14 FcoR genes/individual, and a range of 4 to 28 genes expressed/individual (Extended Data Fig 3).

Similar to the FF loci, the FcoR genes diversify both genetically and somatically, however phylogenetic analysis revealed that the FcoR genes were more complex than the FF genes (Fig. 1d). The FcoR genes grouped into multiple clades, and independent analysis of the ectodomain and TM/intracellular region also showed that these genes diversify via intergenic recombination (Fig. 2c,d). However, the TM/intracellular region of the FcoR genes were more diverse than the FF genes with a higher number of clades in the phylogenetic analysis.

We also found alternative splicing in the extracellular and intracellular regions of FcoR genes (Extended Data Fig. 4b). The most common splicing variant consisted of a deletion upstream of the transmembrane domain, creating a shorter variant of the ectodomain. We also found less common splicing variants, including one that deletes a fragment of the exon that encodes the transmembrane domain, as well as several in the cytoplasmic tail (discussed below).

### Polymorphism of FcoR genes

Polymorphism of the FcoR genes mirrored results from the fester genes. One locus (FcoR7) was highly polymorphic, and we identified 20 allotypes that were 94% identical, with the polymorphisms concentrated in the extracellular region (Extended Data Fig. 2c,d; 5b). Three other polymorphic loci (FcoR2, FcoR3 and FcoR4) were also identified, and were similar to the oligomorphic FF loci (FF12, FF13 and FF28), in that they had 2-3 more divergent alleles (ca. 90% identity, Extended Data Fig. 2c, Table 3). There were also many monomorphic FcoR loci (n>5; Extended Data Table 3). In summary, the patterns seen in the FcoR family are equivalent to the FF family, with the same caveat that almost half of the unique FcoR sequences were present in only one or two individuals.

### FcoRs encode canonical ITIMs, hemITAMs and other tyrosine motifs in the intracellular tails

Every FcoR protein sequence with a transmembrane domain and cytoplasmic tail longer than 100 aa encoded one or more tyrosine-based motifs (Extended Data Fig. 6). Some genes encoded a canonical Immunoreceptor Tyrosine-based Inhibitory Motif (ITIM: I/L/VxYxxI/L/V), others encoded a canonical hemi-Immunoreceptor Tyrosine-based Activation Motif (hemITAM: [E/D][E/D][E/D]xYxxL), and one had an Immunoreceptor Tyrosine-Switch Motif (ITSM: TxYxxI)^29^. Almost all the sequences encoded a tyrosine core motif (YxxI/L/V)^29^. About half of the core motifs had an acidic amino acid at −2 position, and a valine at −1 position ([E/D]VYxxI/L/V), while the other half had no conservation prior to the tyrosine. Several of the sequences also encoded predicted SH2 (Src Homology 2) motifs. Many of the loci encoded multiple motifs, although there was no clear pattern of pairings nor spacing between them (Extended Data Fig. 6).

Two of the polymorphic loci (FcoR3 and FcoR4) encoded alleles that differed in the cytoplasmic tail and swapped the ITIM and hemITAM domains in nearly the same pattern (Fig. 3,). For FcoR4, one of the alleles (FcoR4B) encoded a canonical ITIM motif (LAYAIV) and a partial hemITAM motif (PDDVYAIL), while the other allele (FcoR4A) had a substitution on the ITIM motif (**F**AYAIV), and encoded a canonical hemITAM (EDDVYAIL) (Fig. 3). We also found an alternative splice of FcoR4A in multiple individuals which retained intron 12. This new sequence encoded an ITIM domain (VIYMTI) and excluded the hemITAM (Fig. 3a). Thus, there are activating and inhibitory alleles of the same locus, and these can also be switched by alternative splicing.

**Fig. 3.**
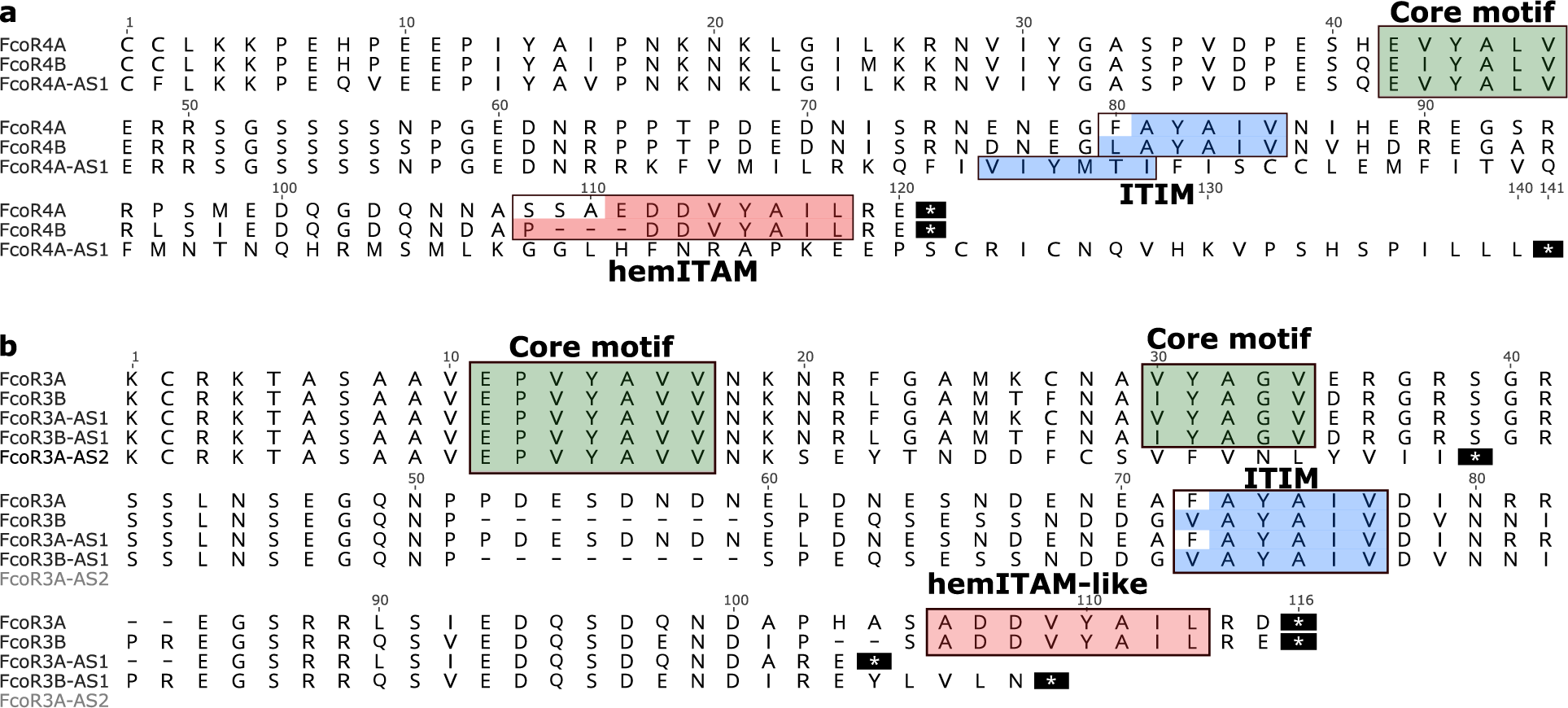
Individual FcoR proteins can switch signaling motifs. **a,** Alignments of the cytoplasmic tail of the FcoR4 alleles (A and B) and alternative splice variant (AS) of 4A. **b,** Cytoplasmic tails of FcoR3 alleles (A and B) and three alternative splice variants (AS). ITIM, hemITAM and tyrosine core motifs are highlighted in blue, red and green, respectively. AS-alternative splice variant.

FcoR3 had an equivalent allelic pattern but was even more diverse, as multiple alternative splice variants were found that modified the signaling motif repertoire (Fig. 3b). Together the data suggests that FcoR can be classified into activating, inhibitory and neutral receptors, and moreover genes can switch motifs via both allelic polymorphism and alternative splicing. Finally, every individual expressed one or more FcoR genes with an ITIM motif, one or more with a hemITAM motif, and one or more with a tyrosine core motif (Extended Data Table 3).

### Co-expression and physical mapping of the FF and FcoR loci

Interestingly, in over half the cases the expression of each FF locus was accompanied by a cognate FcoR locus (Extended Data Fig. 7 and Table 3). We found 14 pairs where loci of the two gene families were co-expressed over 80% of the time in 3 or more individuals, and for five of those the correlation was absolute (Extended Data Fig. 7, Table 3). However, it was the pairings themselves that were striking. Three of the polymorphic FF loci were expressed as pairs with three of the polymorphic FcoR loci, including the two most polymorphic family members (FF1 and FcoR7); as well as two of the less polymorphic family members (FF12 and FcoR3; FF13 and FcoR4). All the alleles of FcoR7 encode an ITIM motif, while both FcoR3 and FcoR4 have alleles with an ITIM motif (Fig.3, Extended Data Fig. 6). The pairing of the only highly polymorphic loci of each family-FF1 and FcoR7, is consistent with our previous functional studies, which suggested the FF1 was part of an inhibitory receptor that discriminated between fuhc ligands, as it was required for fusion^25^. In addition, the paring of the other polymorphic FF with FcoR loci encoding an ITIM is consistent with the idea that the polymorphic FF loci would be involved in specificity, and thus be inhibitory receptors in a missing-self recognition event.

In contrast, functional data suggested that the monomorphic FF3 was an activating receptor^26^, and it pairs with monomorphic FcoR1, which encodes a core tyrosine motif in the intracellular tail (Extended Data Fig. 6c, 7b). In the remaining ten pairs both loci are monomorphic and the FcoR partner encodes a core tyrosine motif.

Physical mapping of the FF and FcoR loci revealed that co-expression patterns correlated with genomic linkage of the two genes (Fig. 4 and Extended Data Fig. 7). We initially mapped the new FF and FcoR sequences back to the partial physical map of the fuhc locus, which consisted of seven scaffolds covering two fuhc haplotypes-(fuhc α and β; Fig. 4a); isolated from California^25^. In these haplotypes, we found that FF and FcoR loci are encoded head-to-head in pairs that correlated exactly with co-expression results from multiple individuals (FF1/FcoR7; FF3/FcoR1; FF22/FcoR42) (Fig. 4a, Extended Data Fig. 7, Table 3). In summary, the FF and FcoR genes are encoded in pairs in two haplotypes and also co-expressed in multiple wild-type individuals. Moreover, presence/absence polymorphism in haplotypes is of loci pairs.

**Fig. 4.**
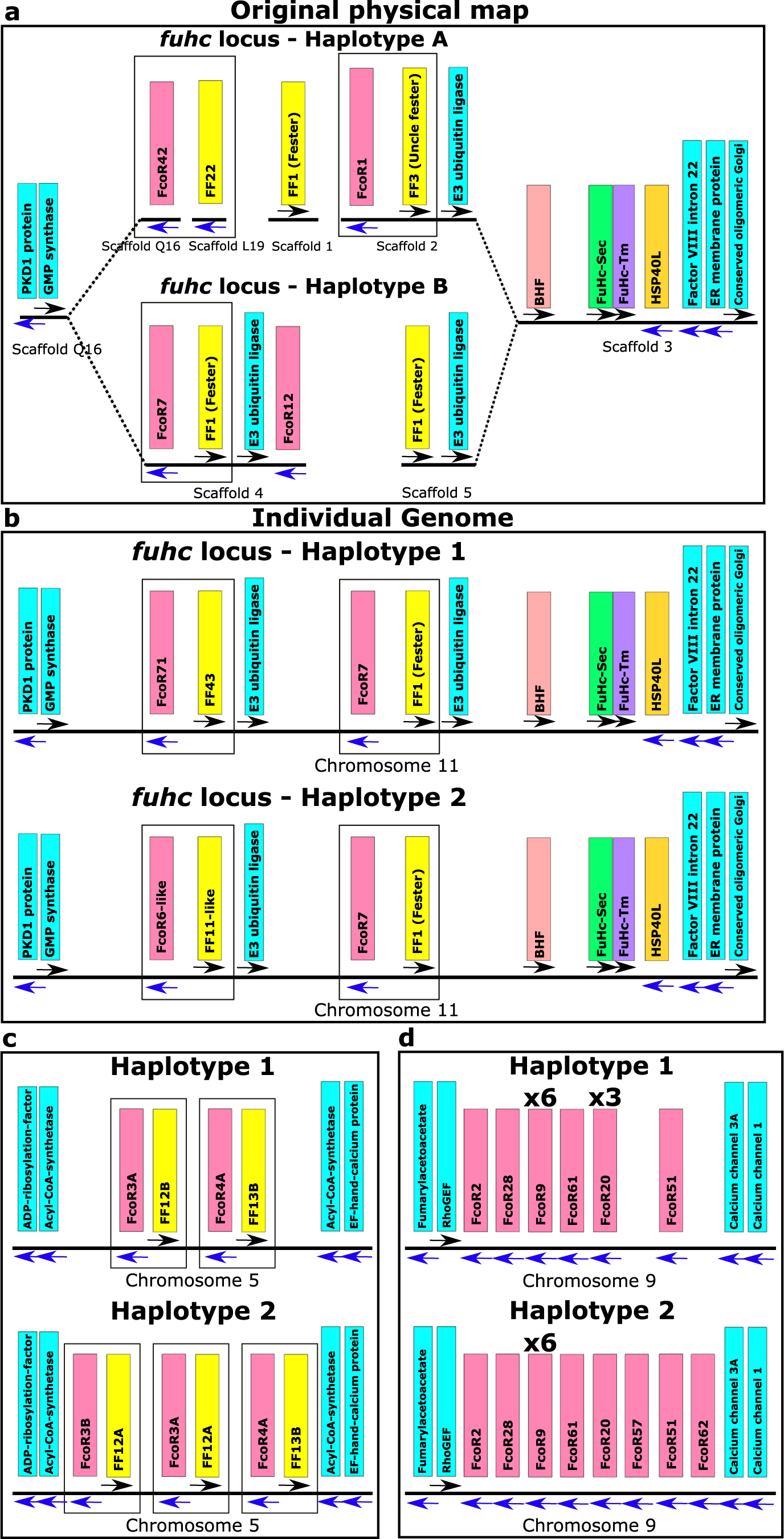
Haplotypic variation of *fester* and *FcoR* genes. **a,** Partial physical maps of the α and β haplotypes of the *fuhc* locus from individuals in California **b,** Comparison of the maternal and paternal haplotype of the fuhc locus (Chromosome 11) from a single individual from the Mediterranean. **c,** Comparison of two haplotypes encoding FF/FcoR pairs on chromosome 5 from the same individual. **d,** Comparison of two haplotypes of the FcoR cluster found on chromosome 9. *fester* and *FcoR* genes are highlighted in yellow and pink, respectively. Framework genes are highlighted in blue.

Next, we analyzed the complete genome sequence of a single individual isolated from the Mediterranean coast of France^30^. In this assembly, both haplotypes of each of the 16 chromosomes were recovered, allowing us to physically map the maternal and paternal chromosomes of this individual independently. Two FF/FcoR clusters are found in the genome, one within the fuhc locus on chromosome 11, and a separate cluster on chromosome 5 (Fig. 4b,c). The physical location of the FF and FcoR loci correlate exactly with the phylogenetic results-(Fig. 1,2) as intergenic recombination would be restricted to genes on the same chromosome. Moreover, despite the geographic distance, three of the FF/FcoR pairs found in individual transcriptomes from California are also encoded in the genome of the Mediterranean individual (FF1/FcoR7; FF12/FcoR3 and FF13/FcoR4), suggesting that the pairings are conserved, and this is notable as these are the three polymorphic pairs of genes (Fig. 4b,c and Extended Data Fig. 9, Table 3).

In addition, a third FcoR cluster is found on chromosome 9 (Fig. 4d). This region is also highly polymorphic, encoding over 13 FcoR genes over an ca. 1.2 Mb region. We detected presence/absence polymorphism between these two haplotypes, and this cluster is larger and more complex than those on the other two chromosomes. The presence of three FcoR clusters is also consistent with the phylogenetic analysis, which had split the FcoR sequences into clades that correspond with their chromosomal location (Fig. 1,2).

Next, we compared the sequenced genomic haplotypes of the fuhc on chromosome 11 (Fig. 4,5). This confirmed our previous results-the fuhc locus is embedded within two groups of conserved framework genes and consists of two parts. On one side, a 120Kb region encodes the candidate fuhc ligand genes: *bhf*, *fuhc-sec*, *fuhc-tm*, and *hsp40l*, (Fig. 4, 5A)^17^, which are conserved between individuals. In contrast, the region spanning from *bhf* to *GMP synthase* genes encodes the highly divergent FF/FcoR haplotypes.

**Fig. 5.**
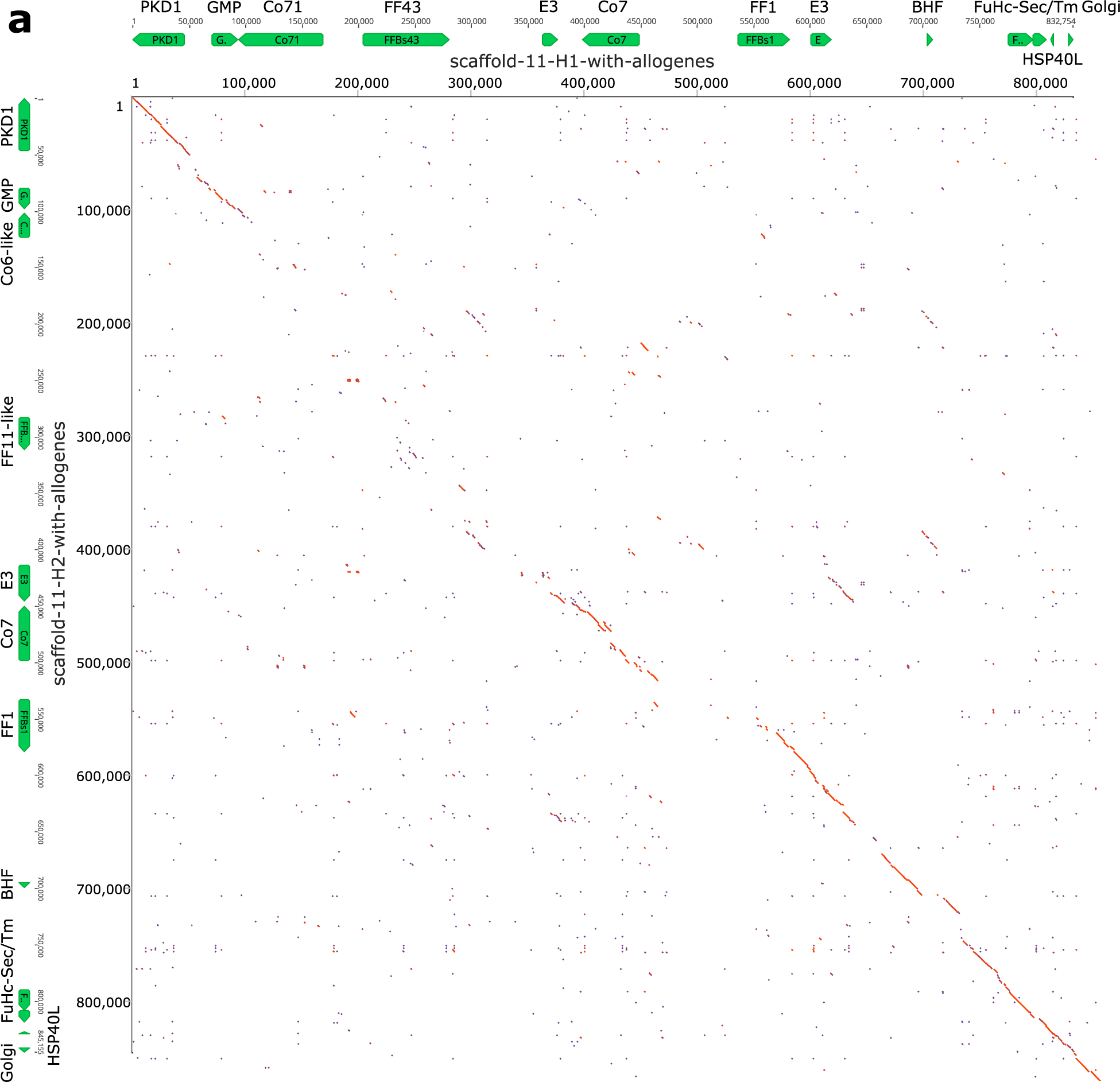
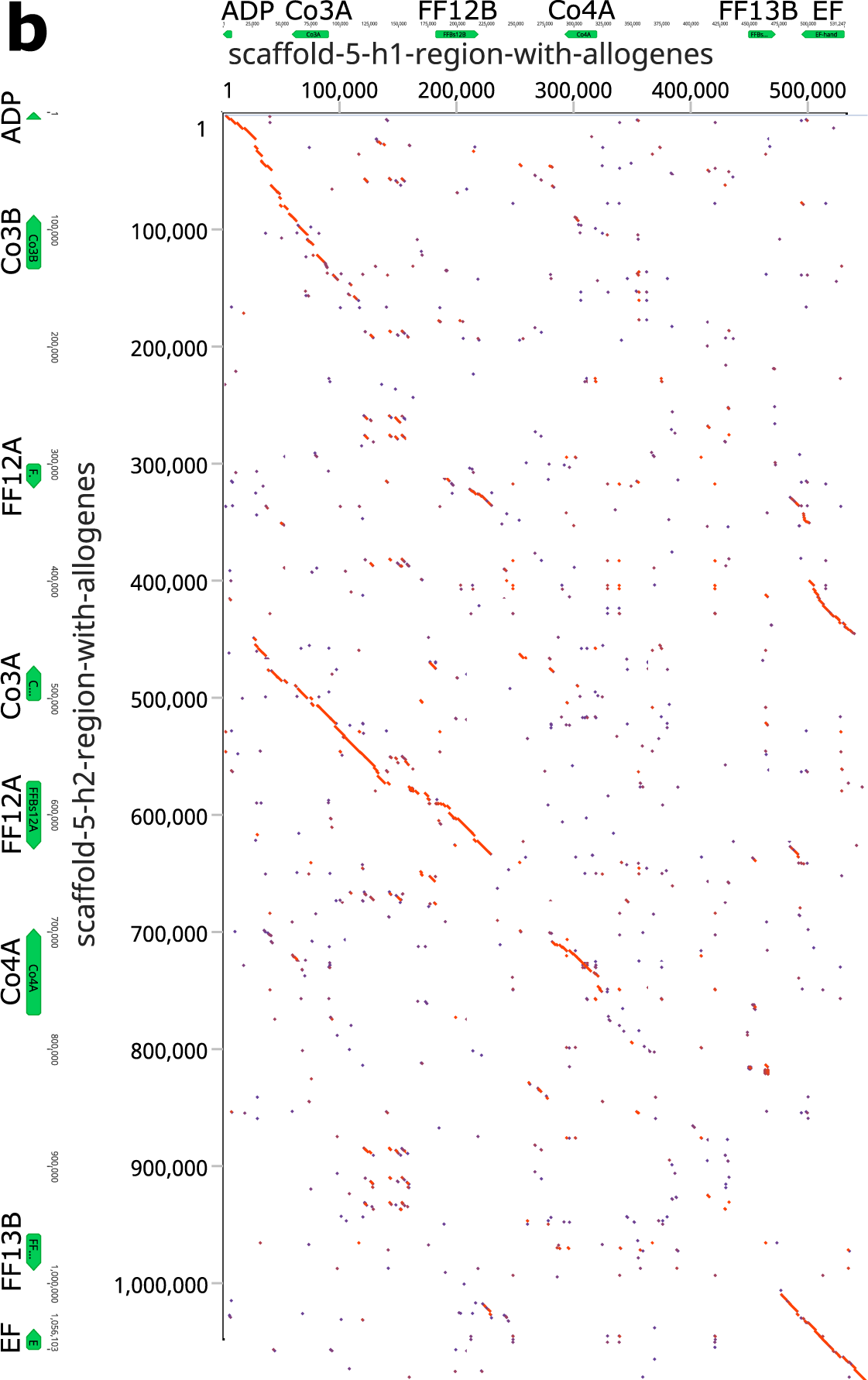
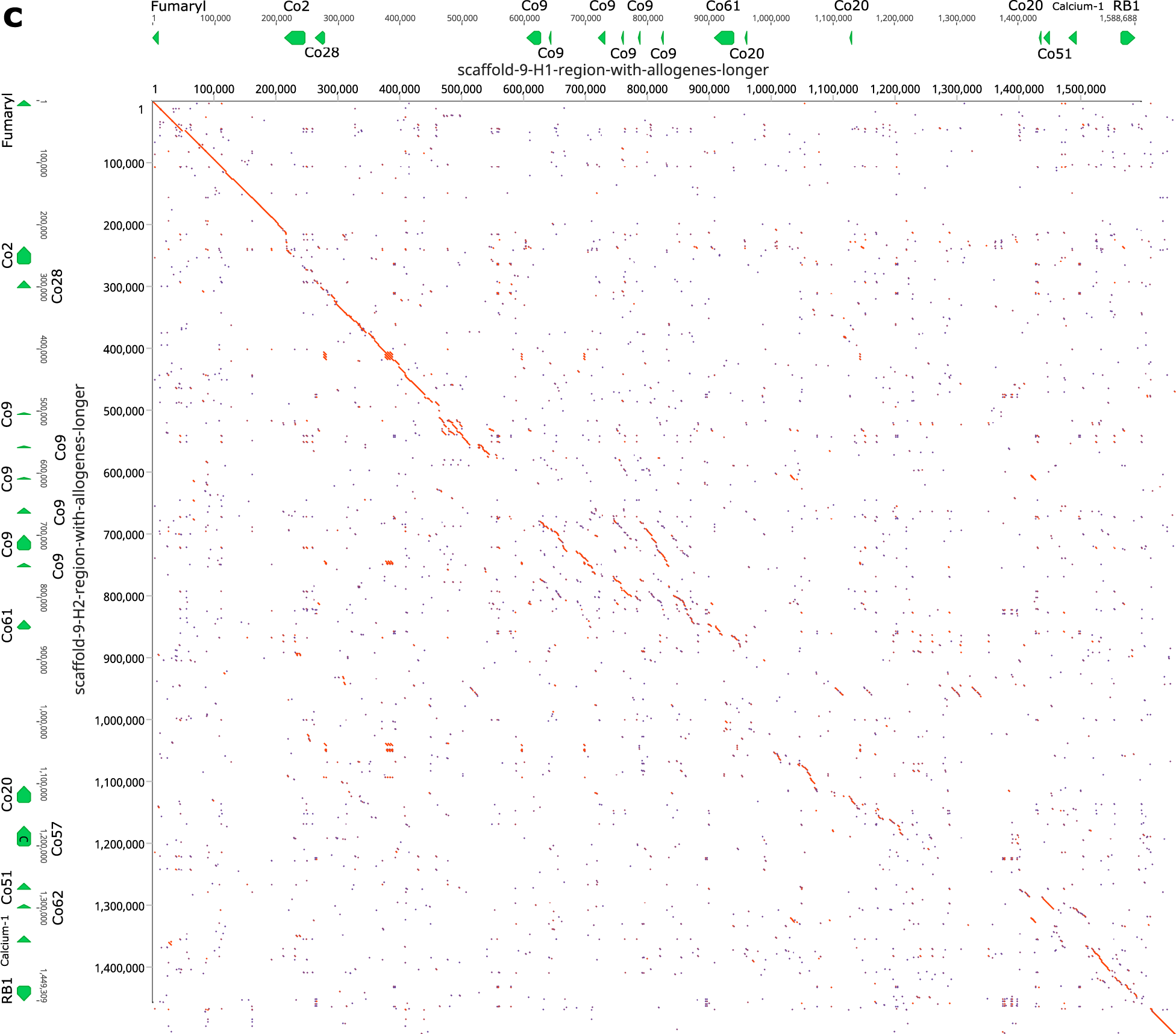
Comparison of maternal ad paternal haplotypes the three allorecognition loci. Dot plot comparison of the two haplotypes from **a,** Chromosome 11. **b,** Chromosome 5. **c,** Chromosome 9. The allorecognition and framework genes are indicated for each haplotype.

Comparison of the FF/FcoR cluster on chromosome 5 and the FcoR cluster on chromosome 9 tell the same story (Fig. 5b,c). The polymorphic receptor clusters are bounded by conserved framework genes, within which these receptors are evolving rapidly by birth and death evolution^31^, with little homology outside the genes themselves. Finally, both the genetic and physical mapping suggests that the majority of diversification is of the FF/FcoR pairs, but is not absolute, as transcriptomic analysis suggests there are cases where one partner is expressed alone (Extended Data Table 3).

### Genotype specific expression of the FF and FcoR genes

In our initial studies we found genotype specific expression and alternative splice patterns of both FF1 and FF3, which we hypothesized were a result of a genotype specific calibration of these two signals that ensures specificity^25,32^. We next surveyed expression patterns of the new FF and FcoR family members in ampullae isolated from different genotypes over multiple time points and found that the new loci were also differentially expressed in stable, genotype specific patterns, regardless of their chromosomal location (Extended Data Fig. 7, 8), consistent with our previous results.

### Botryllus vasculature expresses orthologs of ITAM and ITIM signal transduction proteins

Finally, to determine if the cells involved in allorecognition co-express proteins involved in allorecognition and ITIM and ITAM signal transduction, we characterized gene expression in FACS isolated vascular cells^32,33^. All candidate allorecognition genes in the fuhc locus were expressed in these cells, along with homologs of nearly every signal transduction protein used in ITAM and ITIM signaling in TCR and NK cells, as well as the transcription factors NFAT and NFκB. Vascular cells also express multiple homologs of receptor-type protein phosphatases (Extended Data Fig. 10).

Interestingly, no ascidian genomes encode homologs of DAP12, DAP10, LAT or SLP-76 proteins^30,34^. In summary, vascular cells express all allorecognition proteins as well as all signal transduction and most of the adaptors and transcription factors involved in ITAM and ITIM signaling in T- and NK cells.

## Discussion

Allorecognition in *B. schlosseri* utilizes innate recognition mechanisms to discriminate between alleles of a highly polymorphic ligand. Natural populations have hundreds of transplantation specificities, which means this missing-self recognition system can pinpoint a self-allele from up to a thousand competing alleles. But how this level of specificity is achieved is not understood. In previous studies we had shown that, similar to NK cells, the allorecognition response in Botryllus was not a mutually exclusive outcome, but rather due to the integration of two independent signals, one generated by the binding of fester (FF1), and the other by the binding of uncle fester (FF3), and further that each signal could be manipulated in vivo^25,26^. This suggested that the allorecognition response consisted of a species-specific initiation of a rejection response mediated by a non-polymorphic receptor, which could be overridden by recognition of a self-fuhc allele by a polymorphic receptor. While there were no signaling domains on either protein, we had already found that proteins encoding cytoplasmic ITIM and ITAM domains, as well as all associated signal transduction molecules used in mammals were present in ascidian and other invertebrate genomes^16,34^, so hypothesized that signaling was via pairing with adaptor molecules containing the activating and inhibitory motifs.

Here we find that the fester locus is much more diverse than previously described, with over 37 loci encoded in diverse haplotypes on two chromosomes. These analyses also revealed the presence of an even more diverse gene family, the fester co-receptor (FcoR) genes. The FcoR are type I TM proteins that encode one or more intracellular tyrosine motifs, including canonical ITIM, hemITAM, and two types of tyrosine core motifs. Several also encode SH2 adaptor domains. The FcoR genes are even more diverse than the FF genes: thus far we have identified 53 FcoR loci, and these are encoded in clusters on three chromosomes. On two of those chromosomes (11 and 5) the FF and FcoR are encoded in pairs, while a third cluster on chromosome 9 encodes only FcoR genes (Fig. 4). These data show a diversity of innate receptors as well as linking the Botryllus allorecognition response to canonical immune signaling mechanisms.

While mammalian innate allorecognition receptors share a common polymorphic genomic organization and functional partitioning, ^13,18,35^, the complexity of the Botryllus allorecognition system is unparalleled. The presence of two diverse gene families, their organization into pairs, and localization on multiple chromosomes are unique, and more in line with the increased specificity of Botryllus allorecognition. This redundancy may allow for both maintenance of some interactions via linkage to the fuhc ligand, while the unlinked clusters may allow for independent and rapid evolvability of those loci.

Allorecognition in Botryllus mediates the natural transplantation of germline stem cells between individuals ^36^. The transplant of stem cells is thought to be an altruistic interaction ^37^, but in some genotypes these stem cells can be parasitic, and once transplanted will replace the germline of the other individual ^36^. This is a strong selective force to diversify the fuhc locus, resulting in extraordinary polymorphism that prevents parasitic genotypes from sweeping through a population ^38^. Diversification requires both changes in the ligand, and the ability to correctly detect those changes. Interestingly, the two sides of the locus are evolving using different mechanisms. The candidate ligand genes at the 3’ region of the fuhc locus are linked in a conserved haplotype in all genotypes examined thus far, and diversifying by nucleotide substitutions and small intragenic exchanges ^39–41^. In contrast, the *fester* side at the 5’ region is evolving by birth and death evolution of the loci driven mainly by duplication and recombination (Fig. 2) ^25,28^. We hypothesize that an upper limit on the discriminatory ability carried out by germline encoded proteins would restrain evolution of the ligand, resulting in different mechanisms diversifying each side of the locus.

If the birth and death evolution is a response to the selective pressure to co-evolve with a rapidly evolving ligand^42^, it was surprising to find that out of >35 fester and >50 FcoR newly identified loci, none are as highly polymorphic as FF1 or FcoR7 (Extended Data Fig. 5). Previous results linked function to polymorphism: FF3 was monomorphic and responsible for species-specific activation, while the polymorphic FF1 was an inhibitory receptor discriminating fuhc polymorphisms ^25,26^. Results here are consistent-polymorphic FF are paired with polymorphic FcoR that encode an ITIM, while the non-polymorphic loci are also paired, and the partner FcoR encodes a hemITAM or core motif. In addition, comparing individual haplotypes from California and the Mediterranean show that presence/absence polymorphisms are of FF/FcoR pairs, suggesting that selection does not act on individual loci, but on specificity of the pair. Nevertheless, we would have predicted more polymorphism and inhibitory receptors.

What insight do these results give into the mechanisms underlying allorecognition? We had found that FF1 and FF3 were necessary and sufficient for fusion and rejection pathways, respectively-with one interesting exception (discussed below)^25,26^. The reagents used in those studies did not cross react with new FF family members (Fig. 4a, Extended Data Fig. 2), suggesting that the FF/FcoR are either assembled into, or signal as, oligomers. Furthermore, our previous hypothesis of a single activating and inhibitory pathway was clearly oversimplified ^17^. All full-length FcoR (but no FF) loci encode one or more tyrosine motifs (ITIM, ITSM, hemITAM, core motif), and some also encode SH2 domains. Many loci have combinations of motifs. In addition, both alleles and alternative splice variants of FcoR3 and FcoR4 swap signaling domains, which we hypothesize is important for specificity (Fig. 3; discussed below). While there is no known role of the core motif: YxxI/L/V ^43,44^, it could be part of either activating or inhibitory signaling. Alternatively, these motifs are equivalent to several of the phosphorylation sites on LAT and SLP-76, the two vertebrate signal transduction adaptors conspicuously absent in ascidians. This includes the presence of an acidic amino acid upstream of the tyrosine on half of them, a modification which has been shown to change the speed of phosphorylation, and plays a critical role in polymorphic discrimination by the TCR^45^. If allelic recognition is due to oligomerization of the FF and FcoR proteins at the cell/cell interface, some of these proteins may function as adaptors.

The presence of multiple loci coupled to a diversity of signaling motifs-including the potential to switch between putative activating, inhibitory or neutral functional roles, all indicate that complex tuning of ITAM and ITIM signaling may be required for allelic discrimination. We hypothesize this is due to an interplay of cooperative binding of oligomers that occur both in cis and trans ^46–48^, with the end result is that ITAM/ITIM signals are integrated from multiple binding events, compared to an activation threshold, and the predominant signal determines outcome (fusion or rejection).

For both T- and NK-cells, effector specificity and function are not encoded in the genome. Both cells require a cell autonomous quality control process during development that takes randomly generated binding specificities and sets activation thresholds that allow a cell to be self-tolerant but functional^4,9,10^. But how T-or NK cells establish these thresholds remains unclear. Similarly, specificity in Botryllus does not appear to be hardwired: the extreme polymorphism rules out a simple lock and key interaction between complementary genes, and this mechanism is inconsistent with the genetics of the locus ^17^, and results presented here. We hypothesize that specificity is due to an education process which would modify the stoichiometry of the oligomers until specificity was achieved, reminiscent of the rheostat model of NK education, but with more influence by polymorphisms of the histocompatibility ligand ^49,50^. This is consistent with findings that genotype specific expression and alternative splicing patterns of FF1 and FF3 are maintained following multiple cycles of ablation and regeneration of the ampullae (Extended Data Fig. 7,8) ^25,32^.

In mammals, education is a conserved process required for effector function in five unrelated receptor/ligand pairs: KIR-MHC Class 1; Ly49-MHC Class I; CD94/NKG2-HLA E (Qa-1); Nkrp1-clr-b; and Sirpα-CD47. Each uses a missing-self strategy to determine the health of target cells, and in each knockout of the inhibitory ligand (MHC Class I; Clr-b; or CD47), which would predict an autoimmune phenotype, does exactly the opposite-resulting in tolerance^62,63,64,65^. In addition, in NK cells it has been demonstrated that this tolerance is reversible-NK cells that develop in an MHC deficient background are anergic, but regain functionality after being transplanted into a wild-type background, and wild-type NK cells can acquire tolerance when transplanted into a MHC deficient background ^51,52^. Similarly, knockout of the inhibitory receptor *FF1* blocked both fusion and rejection responses from occurring^16^. We had originally hypothesized that FF1 might be a subunit of both activating and inhibitory receptors^25^. However, in retrospect this could also be revealing an evolutionarily conserved characteristic of ITIM/ITAM signal transduction-constant inhibitory signaling is required for maintenance of effector function in mature cells. This dynamic tuning is also present in T-cells ^53^, and may have clinical consequences on the long-term efficacy of checkpoint therapy.

In summary, our findings suggest that the mechanisms that underlie polymorphic discrimination rely on the dynamic range provided by the interaction between ITAM and ITIM signaling pathways. Both thymic and NK education take receptor diversity and create specificity^4,9,10^, and a similar process existed long before the emergence of vertebrates. Conservation of tunable signal transduction pathways allow receptors to evolve freely, and can explain the rapid, recurrent and convergent evolution of missing self-recognition systems ^13,18,35^, as well as the emergence of adaptive immunity ^12^. The results of this study coupled to the unique characteristics of Botryllus allorecognition^20^ provide a potent model to study these fundamental mechanisms.

## Acknowledgments

We thank Greg Stoney for collecting and maintain the animals used here, and the Center for Scientific Computing (CSC) of the University of California, Santa Barbara for sharing its clusters to perform the bioinformatics analysis. This research was supported by the NIH grants GM139649 and OD030520 to AWD, and the ANR (ANR-14-CE02-0019-01) and INSB-DBM EVOCHORE to ST.

## Author Contributions

HRV and AWD conceived the project and wrote the manuscript. JS assisted HRV with genetic analyses, ST, ODT and FF provided genomic, technical and scientific insights. AWD supervised the research. All authors edited the manuscript and approved the final version.

## Materials and methods

### Animal collection and maintenance

*B. schlosseri* individuals were collected in the Santa Barbara Harbor (CA, US, 34°24’11“N 119°41’31”W). Animals were maintained with seawater flow at 18C and fed with different species of algae (*Nannochloropsis* sp., *Tetraselmis* sp., *Isochrysis* sp., *Dunaliella salina*) (Carolina Biological Supply, US) two times per day a described previously^54^.

### Identification of *fester* and *FcoR* genes

We used the *fester* (DQ517888.1) and *uncle fester* (JF806283.1) genes of *B. schlosseri* as queries to tBLASTx search in different *B. schlosseri* transcriptomes publicly available or previously sequenced in our laboratory (Extended Data Table 1). Raw reads were cleaned of adaptors and low-quality sequences with Trim Galore software using the default parameters (https://github.com/FelixKrueger/TrimGalore). Transcriptome assemblies were performed with the Trinity software using the default parameters^55^. Raw reads were deposited in the GenBank under the accession numbers ###. Assembled transcriptomes and gene alignments were deposited on GitHub. To identify *FcoR* genes, the *fester* genes from *B. schlosseri* were initially used as queries in a search (tBLASTx) in the transcriptomes and genomes of this species. Finally, identified *FcoR* genes were used as queries in a search (tBLASTx) for additional *FcoR* genes in the databases of this species. Alignments were performed using the BioEdit software^56^.

### Quantification of *fester* and *FcoR* genes

To quantify the expression of *fester* and *FcoR* genes, raw reads from *B. schlosseri* transcriptomes were mapped to the full-length of *fester* and *FcoR* genes using the kallisto method^57^. Trimmed Mean of M-values (TMM) were used to quantify the expression of *fester* and *FcoR* genes. To estimate the number of *fester* and *FcoR* genes per individual, an average was made between the number of *fester* genes present in the replicates corresponding to each individual.

### Identity matrices of *fester and FcoR* genes and loci/allele predictions

Multiple alignments of *fester* and *FcoR* genes were performed with ClustalW and identity matrices were generated with the BioEdit software ^56^. Data visualization was performed with ggplot2 in RStudio (https://posit.co/). To predict which sequences represented unique loci vs alleles, we used amino acid and nucleotide identity values from the most divergent alleles identified in a previous population genetic study of *fester* as a metric, which analyzed >60 alleles found in populations from both the East and West Coast of the U.S.^28^. We identified sequences that had > 80% identity at the amino acid level (Extended Data Fig. 1). If those sequences also shared >90% nucleotide identity over the entire gene sequence, we classified these as alleles. This seemed to be a conservative threshold for both *FF* and *FcoR*, as the only sequences that had nucleotide alignment over their entire length were >90% identical. No other pairs of sequences had homology over the entire length.

### Amplification of *fester* genes from genomic DNA and mRNA

*fester* genes of *B. schlosseri* were amplified from genomic DNA and mRNA. mRNA sequences of *B. schlosseri fester* genes were aligned with the BioEdit software ^56^, and primers were designed for each *fester* gene. For genomic amplification, the size of the amplified regions was between 100 and 200 bp, which sought the amplification of genomic fragments belonging to single exons. Meanwhile, primers covering the full-length of the *fester* genes (approximately 1000 bp) were designed for amplification from mRNA. A full list of primers used in this study is provided in Extended Data Table 2. Genomic DNA was isolated from whole colonies of *B. schlosseri* with the NucleoSpin Tissue, Mini kit for DNA from cells and tissue (Macherey-Nagel) following the manufacturer’s protocol. Meanwhile, total RNA was isolated for the same individuals with the Monarch® Total RNA Miniprep Kit (New England Biolabs) following the manufacturer’s protocol. cDNA was synthetized by adding 200units M-MuLV Reverse Transcriptase, 1X M-MuLV Reverse Transcriptase Reaction Buffer (New England Biolabs), 5mM DTT, 40U RNaseOUT (Thermo Fisher), incubating at 42°C for 1h, and at 70°C for 15 minutes. Gene amplification was performed with the Classic Taq DNA Polymerase Master Mix (Tonbo Biosciences) and 0.5uM of each primer. Amplification protocol consisted of an initial denaturation at 95°C for 2 minutes and 30 seconds, a second denaturation at 95°C for 20 seconds, an annealing step at 57°C for 20 sec and an elongation step at 72°C for 20 sec, repeating these last three steps 35 times, and a final elongation step at 72°C for 5 minutes. PCR products were analyzed by gel electrophoresis.

### Genomic annotation of *fester and FcoR* genes

*fester* and *FcoR* transcriptomic sequences were used as queries (tBLASTx) to identify the genomic scaffolds of *B. schlosseri* containing these genes. Genomic scaffolds were obtained from^25^ and a draft genome of *B. schlosseri*^30^. Transcripts sequences were then manually aligned with the genomic scaffolds to establish the exon-intron boundaries of the *fester* and *FcoR* genes. Genomic scaffolds were annotated with the GenScan software (http://hollywood.mit.edu/GENSCAN.html)^58^and BLASTp using the non-redundant protein sequences (nr). Additionally, tyrosine-based motifs (SH2 motifs) of FcoR proteins were predicted with the Scansite 4.0 software (https://scansite4.mit.edu/#home)^59^, or manually annotated following reported sequences of ITIM ([S/I/V/L]xYxx[I/V/L]; where x represents any amino acid, hemITAM (three acidic amino acids [D/E] follow by xYxxL), ITSM ([S/T]xYxx[L/I]), and two ‘core’ tyrosine-based (Yxx[L/V/I]; [V/I]Yxx[V/L]) motifs^29^.

### Linkage analysis between *fester* genes and the *fuhc* locus

*fester* genes were amplified from genomic DNA of fuhc scored F_2_ and backcross progeny from defined crosses using five *fuhc* haplotypes (α, β, X, C, D)^25^. The PCR amplification protocol from genomic DNA was the same as that described above. Primers are provided in Extended Data Table 2. PCR products were analyzed by gel electrophoresis.

### Phylogenetic analysis of *fester and FcoR* genes

Alignments of the full-length fester and FcoR proteins of *B. schlosseri* were performed with Muscle software (https://www.ebi.ac.uk/Tools/msa/muscle/)^60^. Phylogenetic trees were generated by the Maximum likelihood method with 10000 bootstrap replicates using the IQ-TREE web server (http://iqtree.cibiv.univie.ac.at/)^61^, and edited with the iTOL software (https://itol.embl.de/upload.cgi)^62^. Additionally, phylogenetic trees with the extracellular and transmembrane/cytoplasmic regions of the fester and FcoR proteins were generated using the same strategy described above. Transmembrane domains of the fester and FcoR proteins were predicted with InterProScan software (https://www.ebi.ac.uk/interpro/search/sequence/)^63^.

### Comparison of genomic regions with *fester and FcoR* genes

Genomic regions with *fester* and *FcoR* genes were *in silico* isolated from two different haplotypes (h1 and h2) of *B. schlosseri*. These genomic regions from chromosomes 5, 9 and 11 were pairwise aligned using Emboss stretcher^64^. Dot plots were generated using Geneious Prime.

### Polymorphism of *fester and FcoR* genes

Amino acid diversity of *fester* and *FcoR* genes was calculated using the diversity value (*d*)^62,65^. *BsFesterCo7* sequences were isolated from different *B. schlosseri* transcriptomes. *FF1* sequences were previously reported^28^. Alignments were performed using the BioEdit software ^56^. Redundant sequences were removed before diversity values were calculated. Amino acid positions with 100% conservation (diversity value=0.05) were excluded in the visualization of the results.

### Identification of NK-cell gene homologs

Human ITAM and ITIM signal transduction proteins were collected from the KEGG database (https://www.genome.jp/kegg/), and used as queries (tBLASTx) to search a publicly available transcriptome database of *B. schlosseri* (http://octopus.obs-vlfr.fr/). Putative homologs were annotated with BLASTp and InterProscan software to determine the identity and domain architecture of these proteins. Expression was characterized in FACS isolated vascular cells as described^18^. *Botryllus* homologs were deposited in GenBank.

## Extended Data

**Extended Data Fig. 1.**
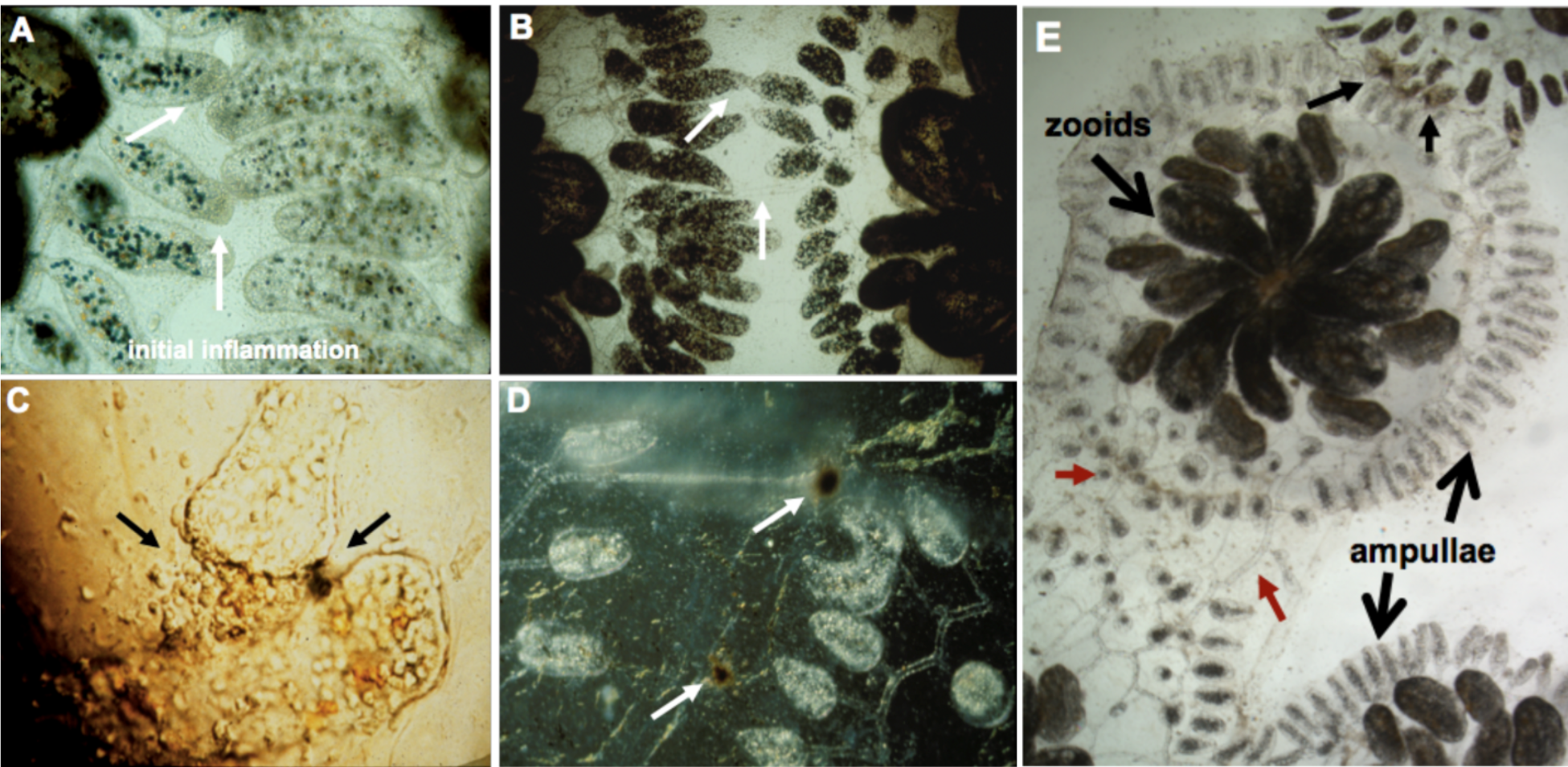
Allorecognition in *Botryllus schlosseri.* **a**, When two colonies grow close together, the ampullae reach out and interact (white arrows), and this will result in one of two outcomes: either the two ampullae will fuse **b**, allowing the circulation of the two colonies to interconnect (white arrows), or they will reject each other (**c,d**). Rejection is a localized inflammatory reaction where blood cells leak from the ampullae. **c**, A close-up of rejecting ampullae showing cell leakage. Once outside the circulation, cells discharge their vacuoles, initiating a prophenoloxidase pathway which eventually forms dark melanin scars, called points of rejection, or POR (**c**, black arrow on right; **d** white arrows). The ampullae then disintegrate (left of POR, top white arrow in **d**), and the colonies no longer interact. The reaction takes ∼24-48h to occur and is controlled by a single highly polymorphic locus called the *fuhc* (for fusion/histocompatibility). Colonies will fuse if they share one or both alleles, and will reject if no alleles are shared. **e**, Lower magnification shows a single individual, consisting of multiple zooids occupying the center. The large extracorporeal vasculature and terminating ampullae are outlined (large arrows). In this experiment, a colony was placed between a compatible partner (bottom) and an incompatible partner (top). As shown, a colony can simultaneously fuse (red arrows) and reject (small black arrows, top). This demonstrates that allorecognition is spatially segregated and occur indpendently at the tips of the ampullae that are in contact.

**Extended Data Fig. 2.**
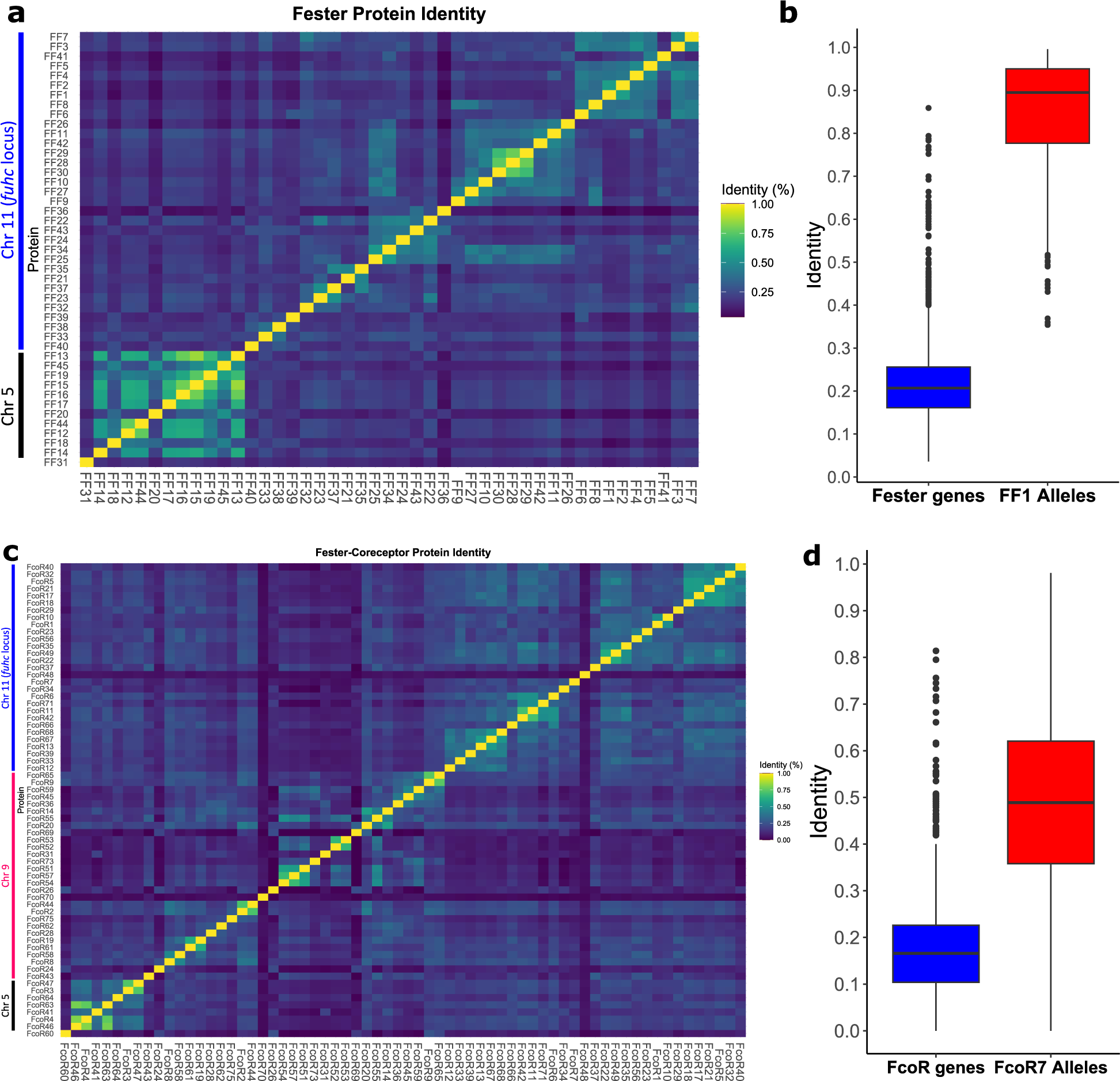
Identity matrix for fester and FcoR genes. **a,** Identity matrix of *fester* genes. **b,** Comparison of identity between the *fester* genes and *FF1* alleles. **c,** Identity matrix of *FcoR* genes. **d,** Comparison of identity between the *FcoR* genes and *FcoR7* alleles. We used the range of amino acid and nucleotide identify from FF1 and FcoR7 alleles as a benchmark to predict alleles vs. loci of the new sequences. If sequences shared >82% amino acid homology and >90% nucleotide homology over the entire gene, they were classified as alleles.

**Extended Data Fig. 3.**
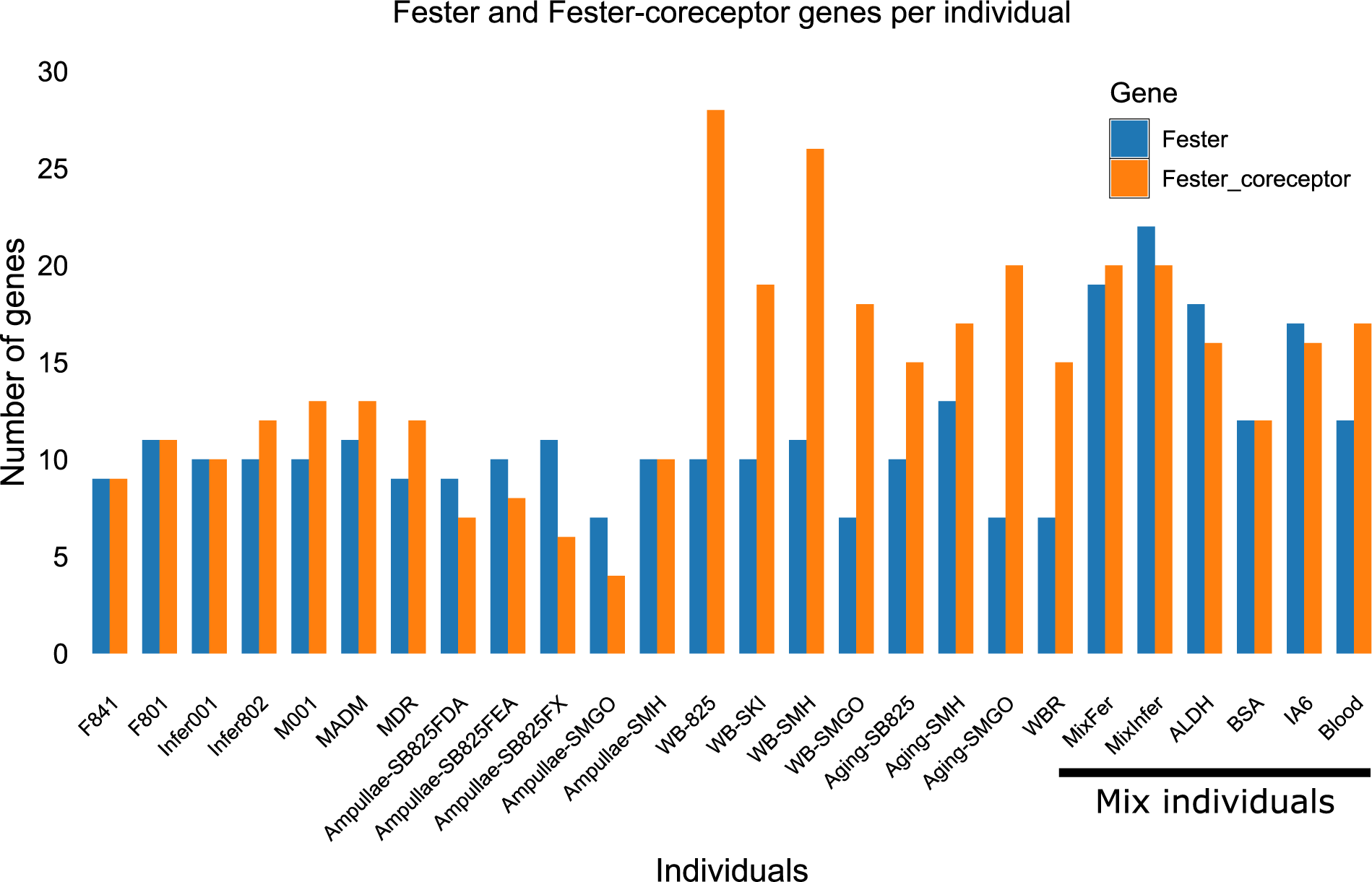
Expression of *fester* and *FcoR* genes. *fester* and *FcoR* genes were quantified in different individuals of *B. schlosseri* using transcriptome assemblies. The last six samples on right contain a mix of individuals.

**Extended Data Fig. 4.**
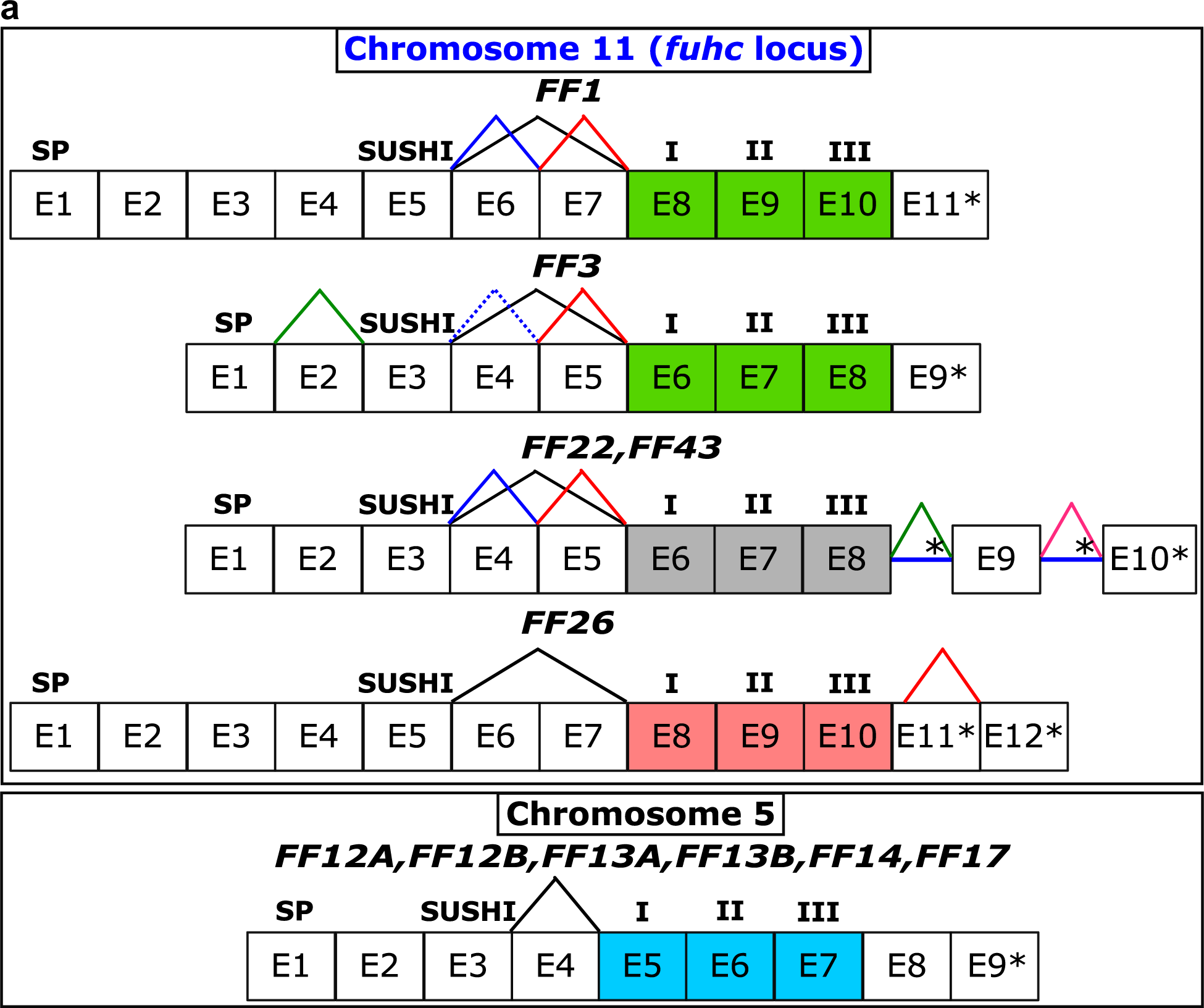
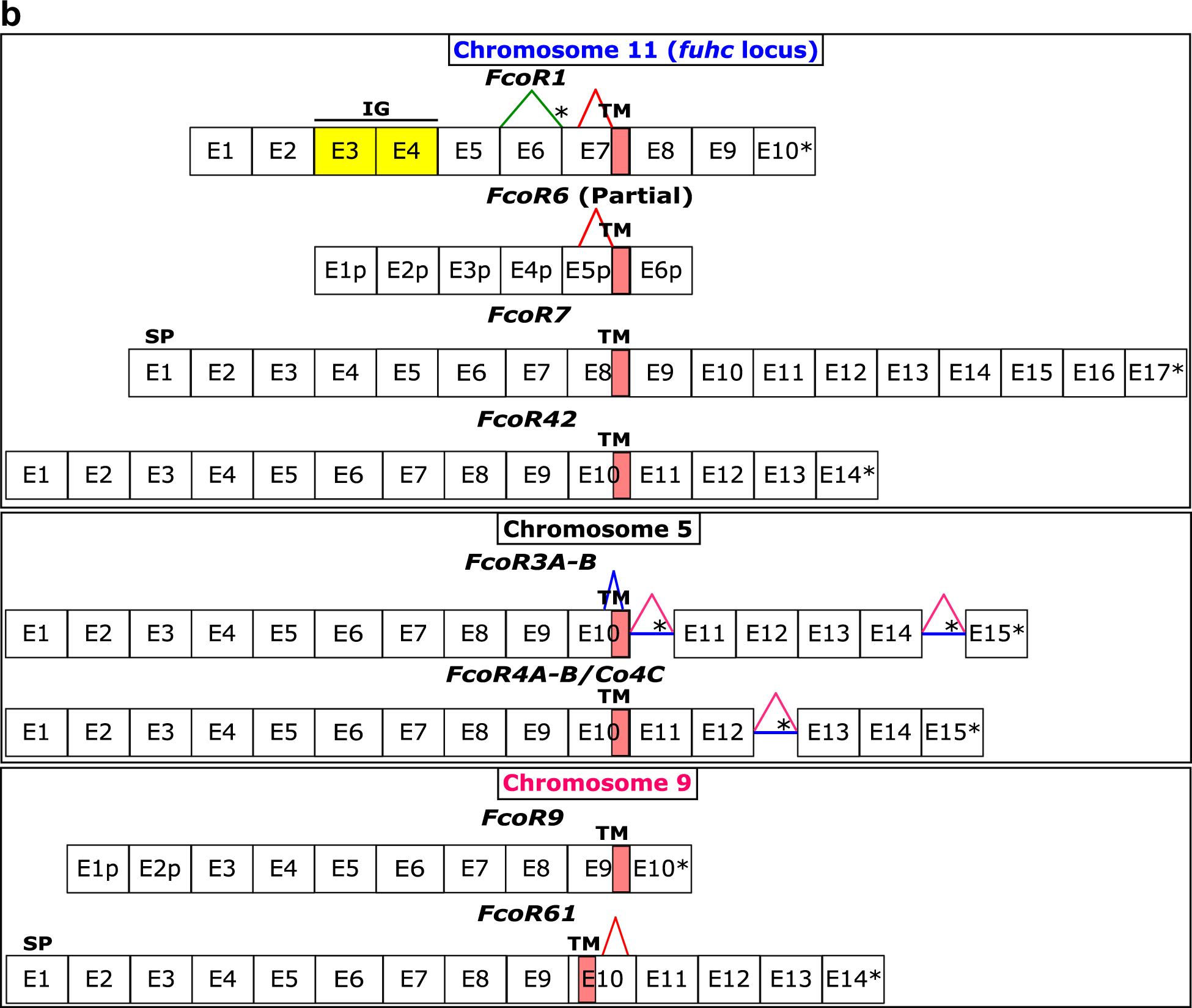
Genomic structure and alternative splicing of *fester* and *FcoR* genes. **a,** *Fester* genes exhibit two regions with alternative splicing. The first region is between the Sushi domain and the first transmembrane domain. The second region is located in the cytoplasmic region. Exons encoding the transmembrane domains are highlighted in colors. **b,** *FcoR* genes exhibit alternative splicing in the extracellular and intracellular regions. Asterisks represent stop codons. Intron retentions are indicated with blue horizontal lines. Signal peptide: SP. I, II, and III: First, second and third transmembrane domains, respectively.

**Extended Data Fig. 5.**
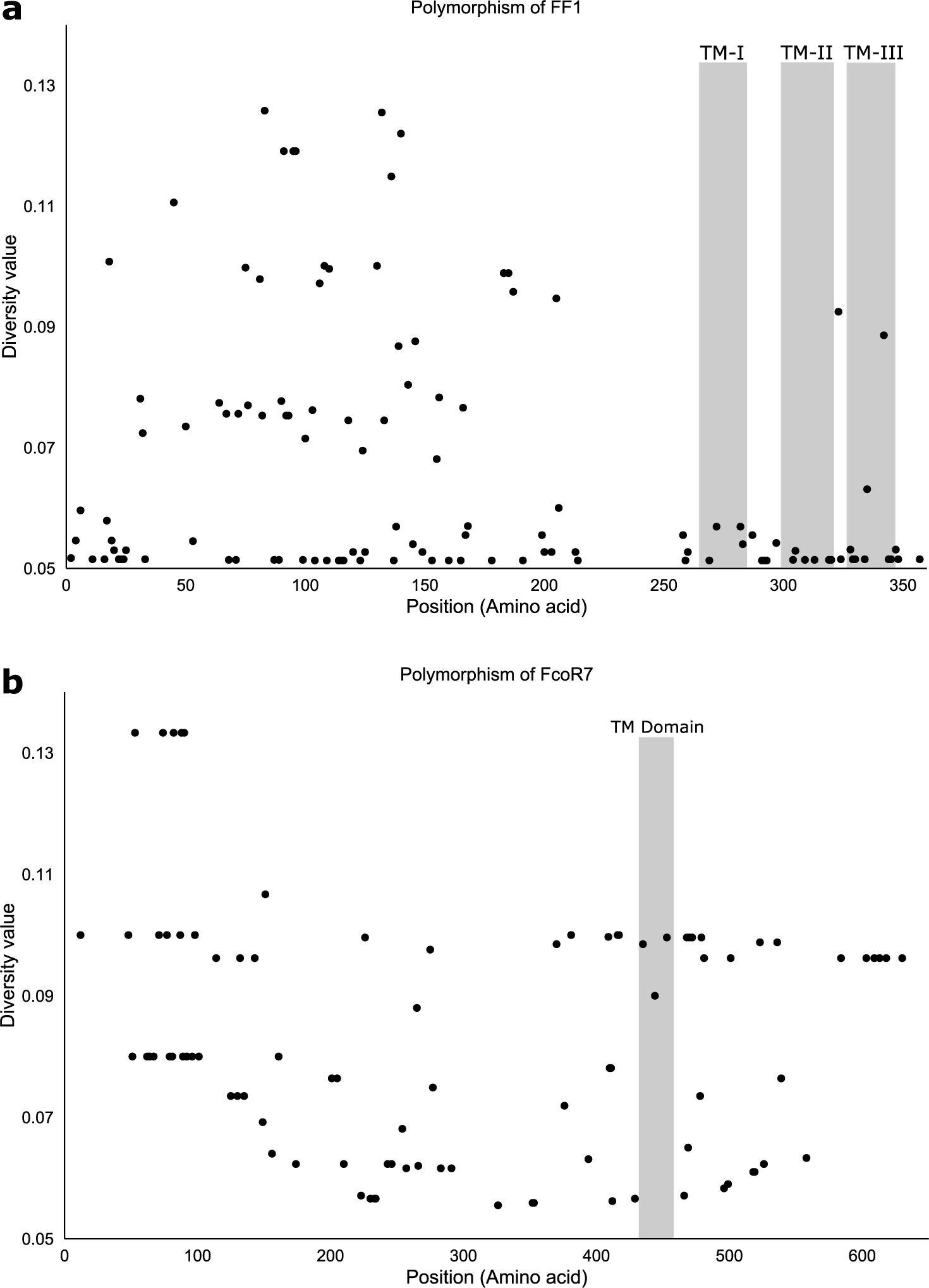
Distribution of polymorphic residues in FF1 and FcoR7 alleles. **a,** 77 allotypes of the FF1 protein and, **b,** Twenty allotypes of the FcoR7 protein were compared using DIVAA. Only variable residues are represented. Amino acid position and diversity value are represented in the X-axis and Y-axis, respectively. A diversity value of 1 for a particular position means that any amino acid is as likely as any other to be present at that position. A diversity value of 0.05 is consistent with 100% conservation at that site (1/20 possible amino acids).TM: Transmembrane domain.

**Extended Data Fig. 6.**
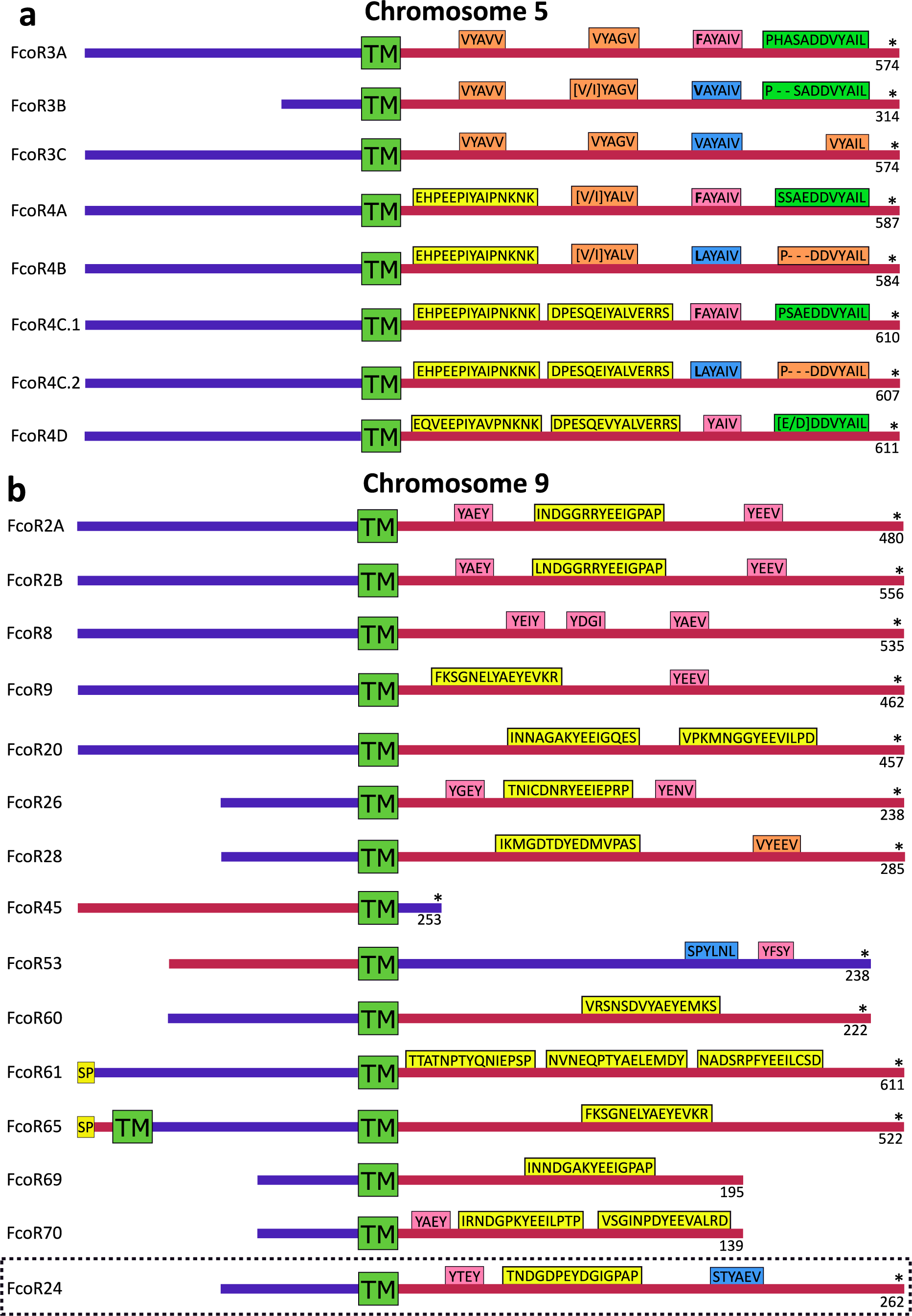
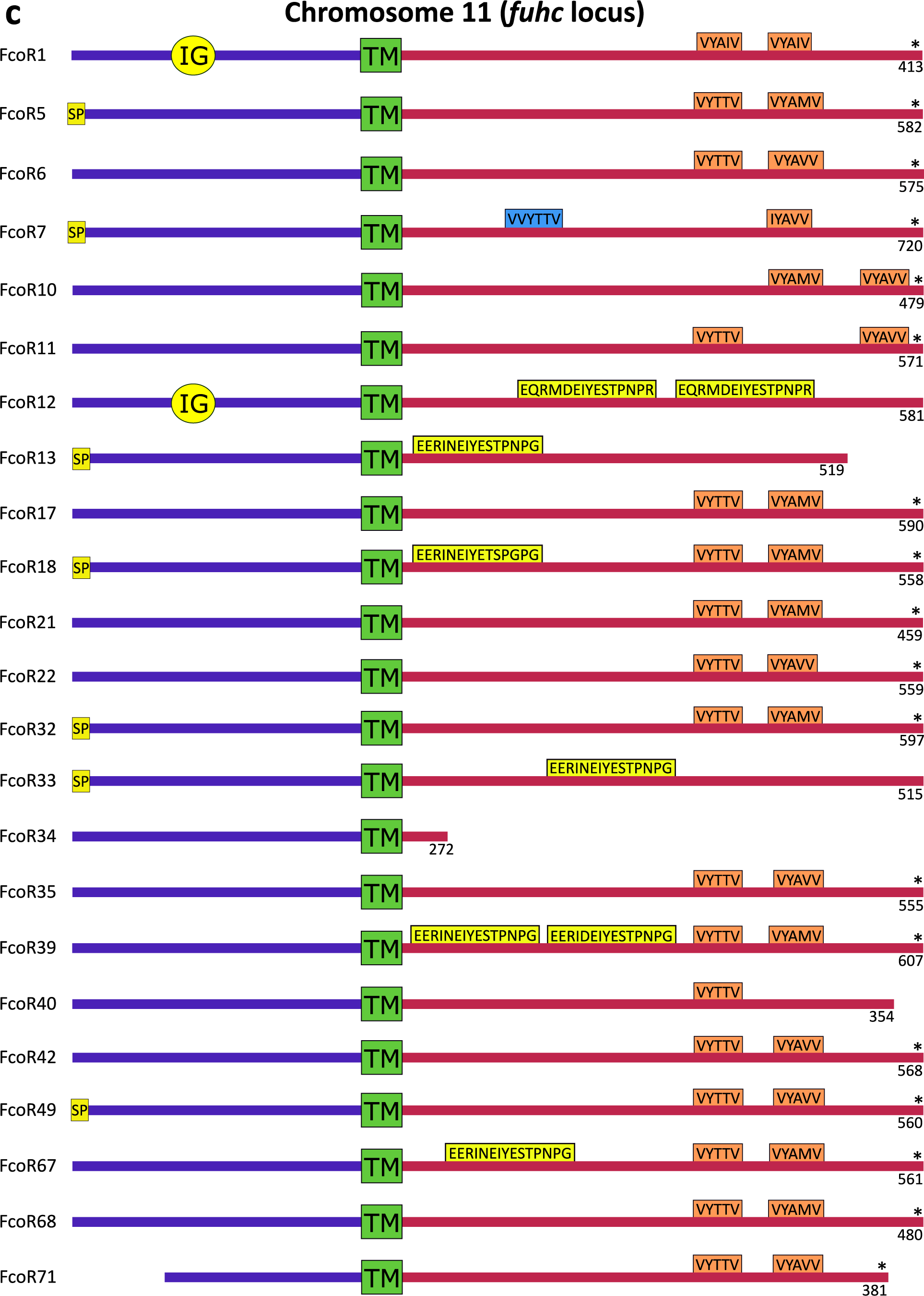
Structure of FcoR proteins. Protein structure of FcoR proteins located on: **a,** chromosome 5. **b,** chromosome 9. **c,** chromosome 11. SP: Signal peptide. IG: Immunoglobulin domain. TM: Transmembrane domain. ITIM, hemITAM, SH2, Tyrosine core-1 (YxxY/L/V/I) and Tyrosine core-2 (V/IYxxV/L) motifs are highlighted in blue, green, yellow, pink and orange colors, respectively. Blue and red lines represent the extracellular and intracellular regions of FcoR proteins,* = stop codon. FcoR proteins without a SP are not likely full-length, as there is no initiator methionine. Partial sequences without a transmembrane domain are not shown.

**Extended Data Fig. 7.**
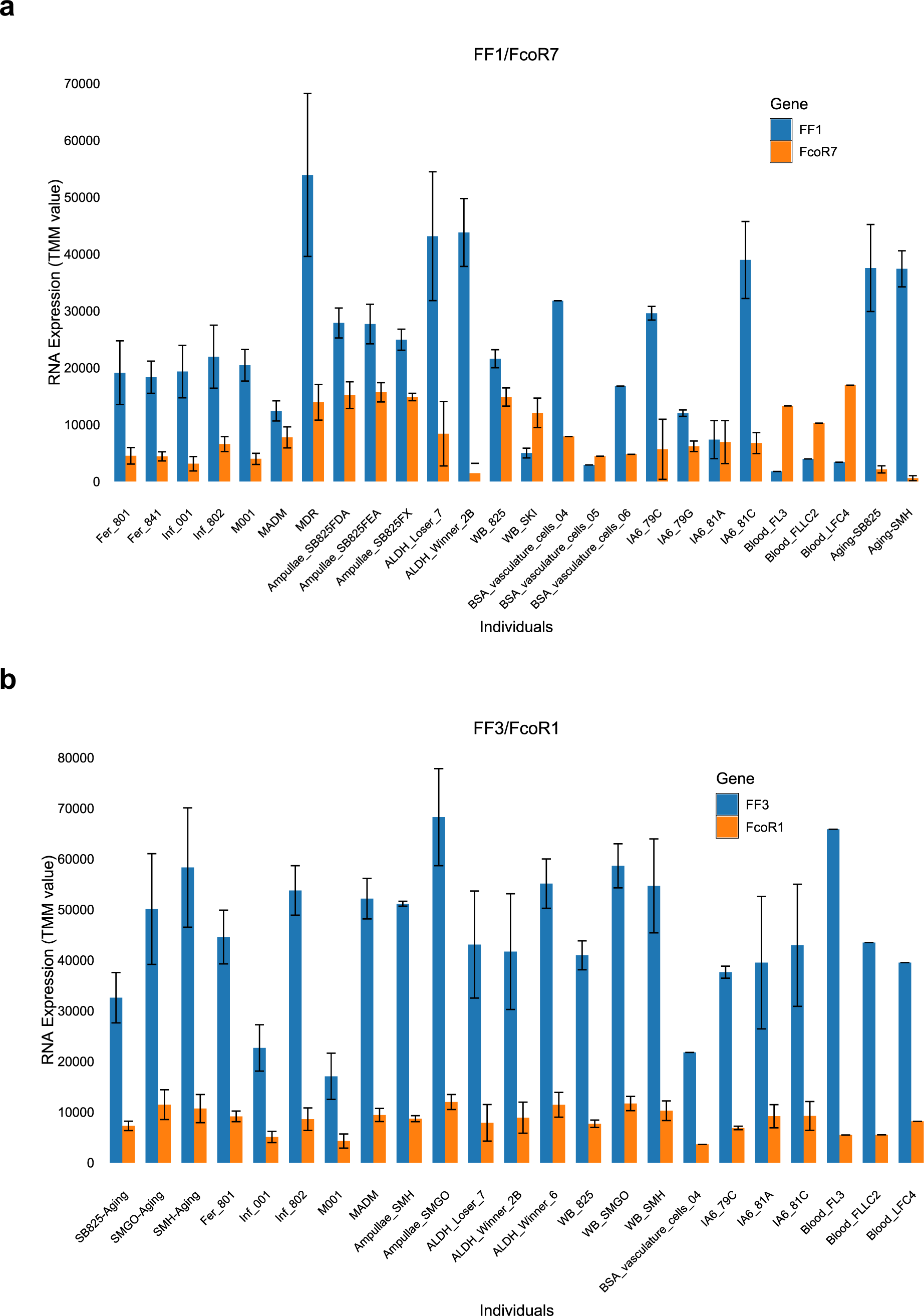
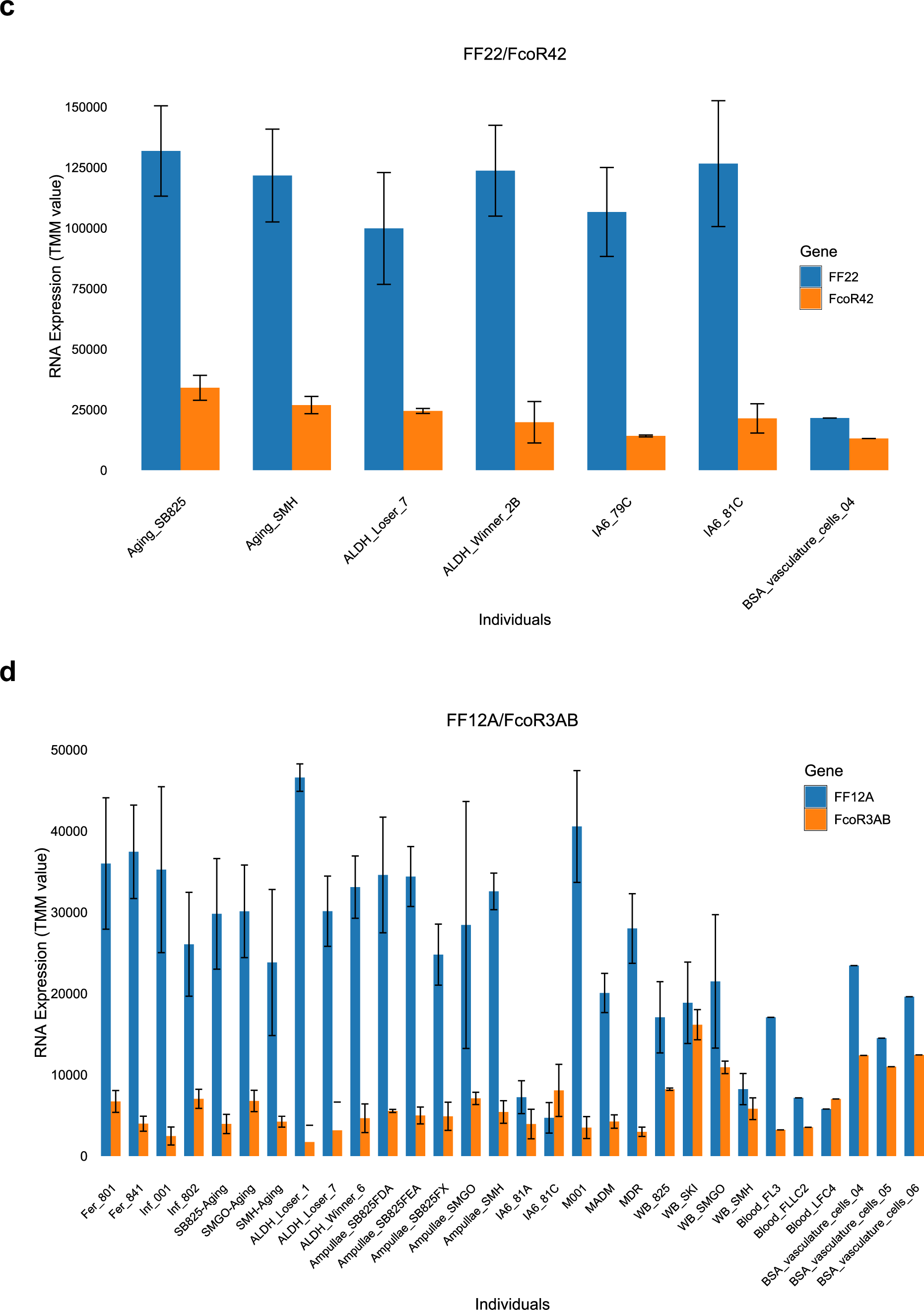
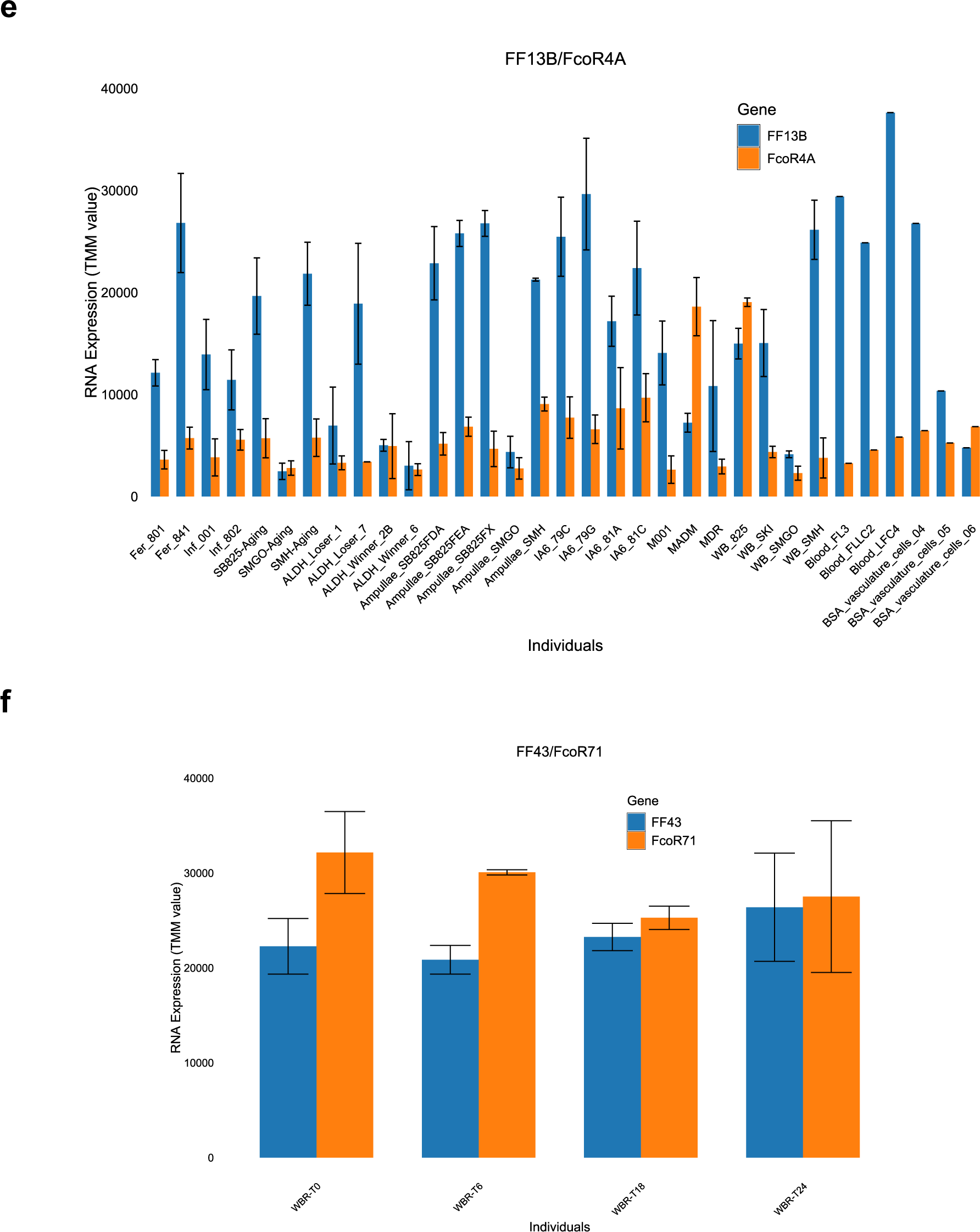
Co-expression of *fester* and *FcoR*. *fester* and *FcoR* gene pairs were quantified in different individuals of *B. schlosseri*. X-axis shows different *Botryllus* individuals, and Y-axis shows gene expression levels (TMM units).

**Extended Data Fig. 8.**
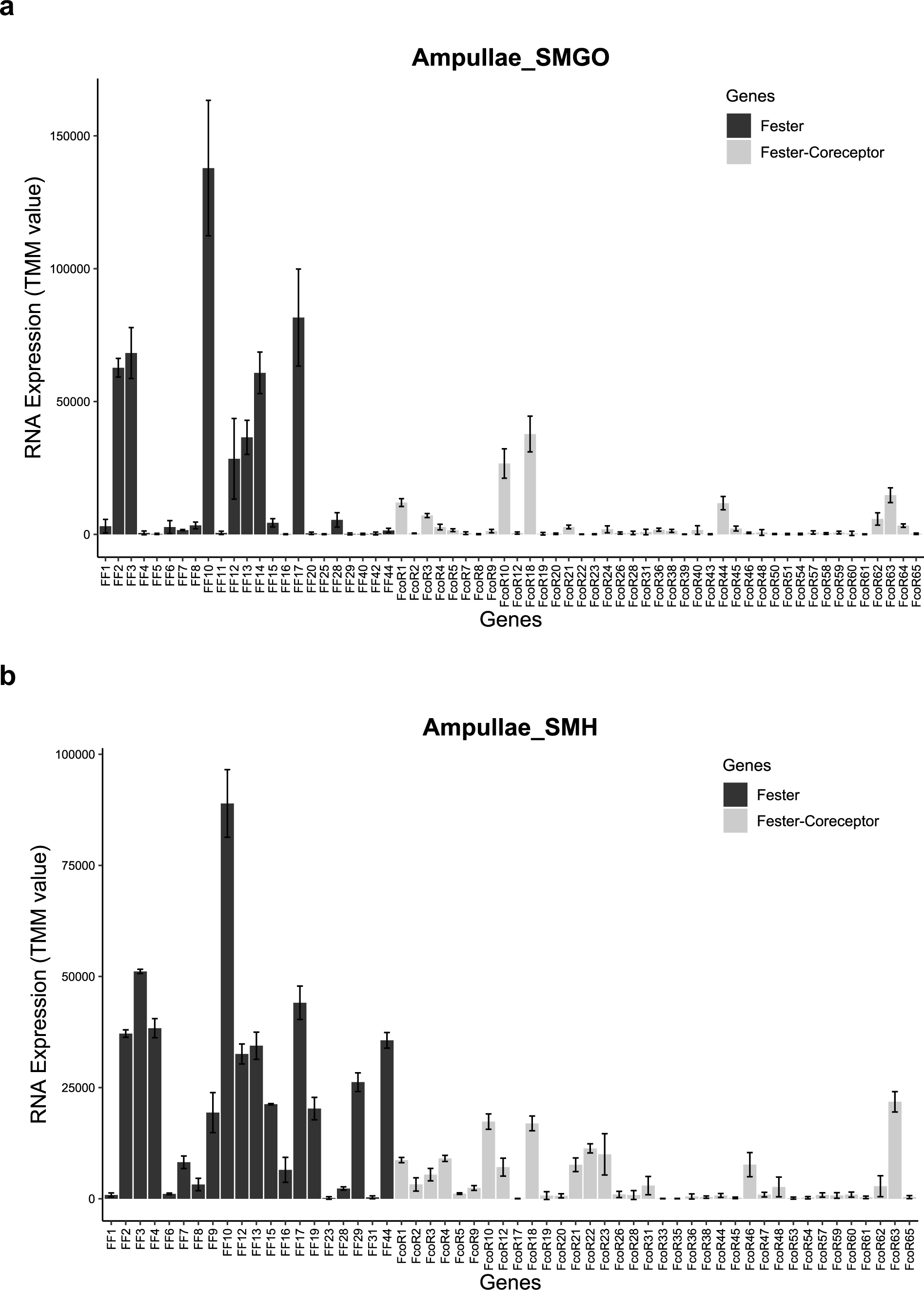
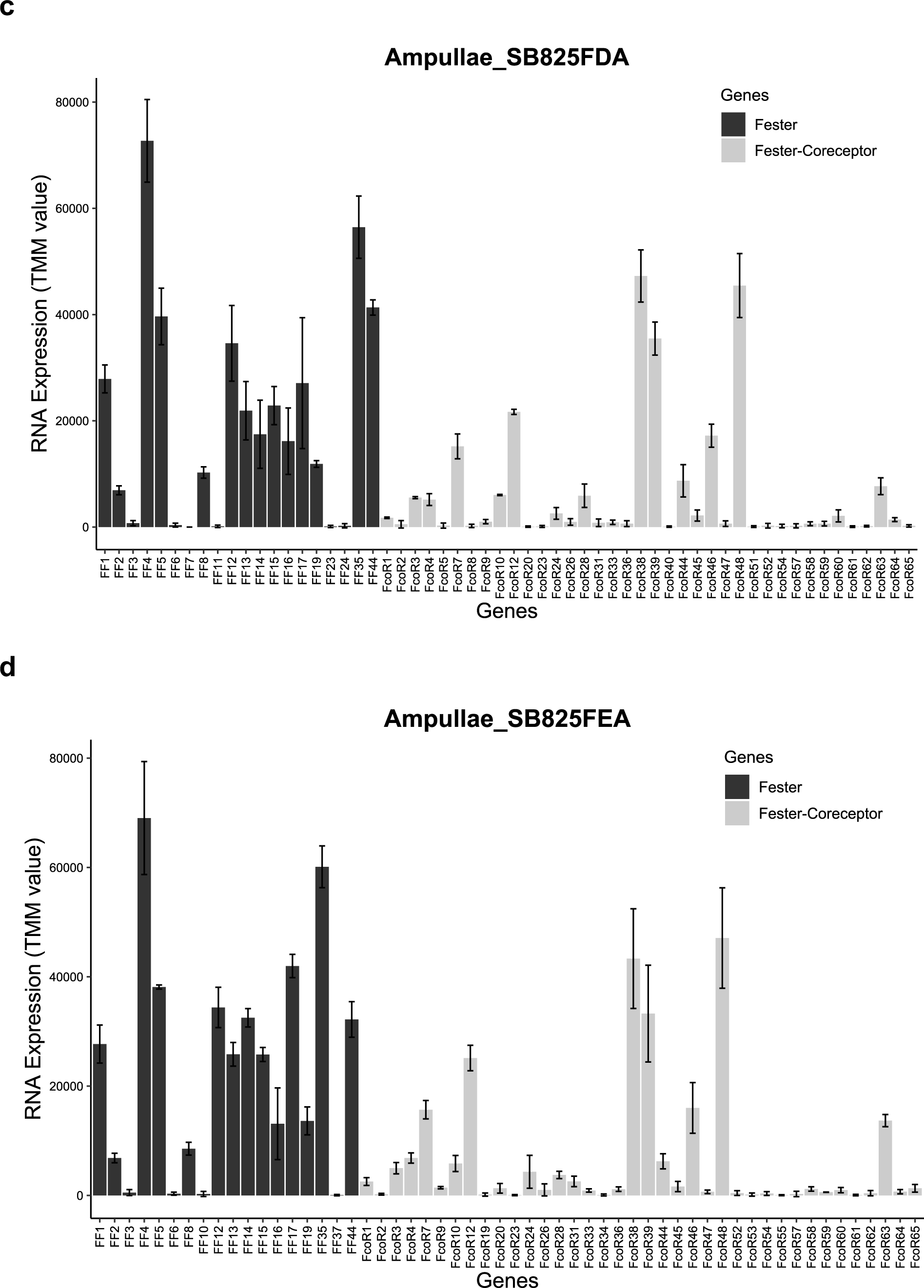
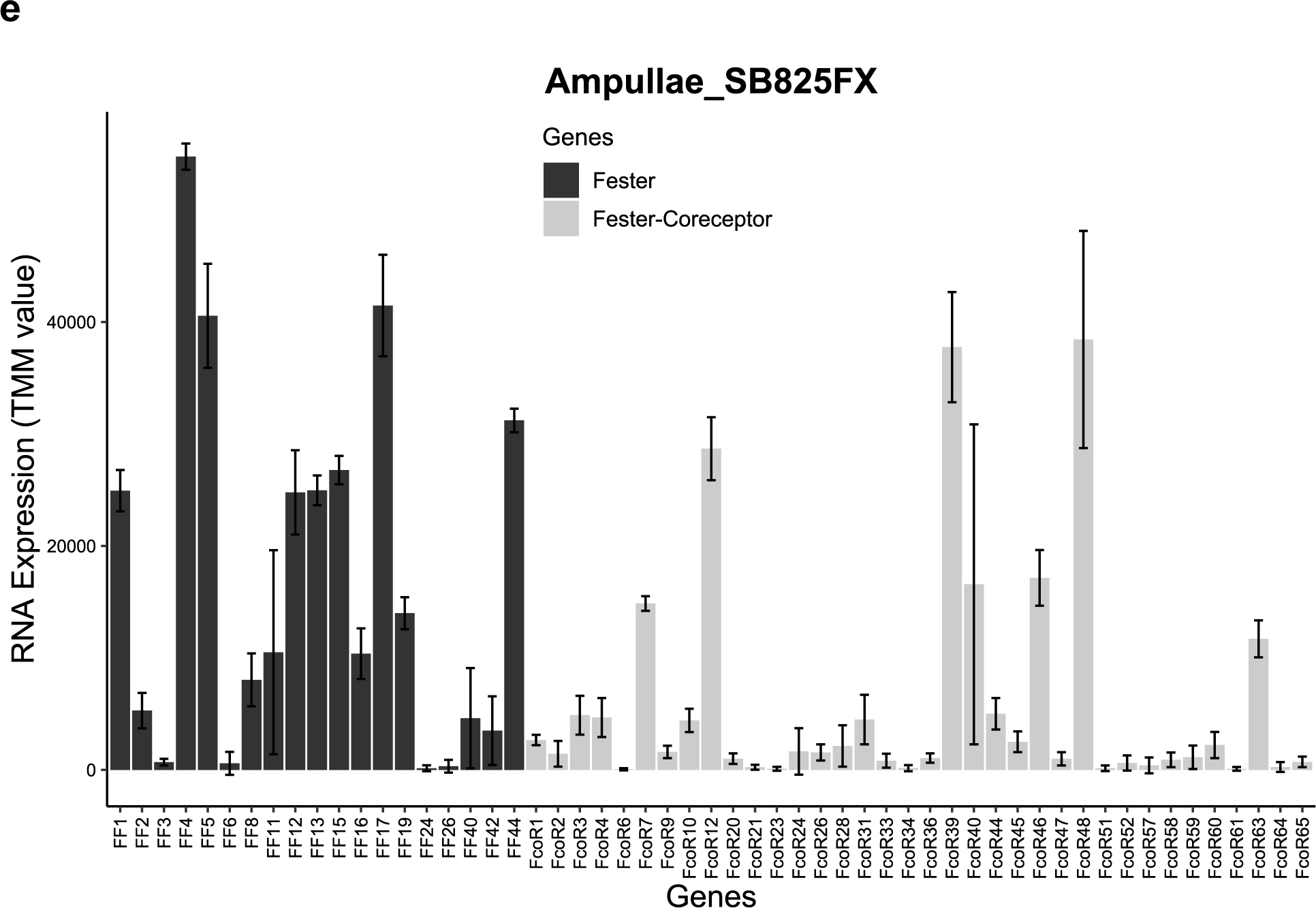
Unique repertoire and stable expression of *fester* and *FcoR* genes in ampullae isolated from 5 individuals. **(a-e).** In each individual, ampullae were surgically removed and collected, the vessels regenerated, and this tissue isolation/regeneration was done three times. mRNA isolated from each sample was sequenced, and the average expression (Y axis, TMM units) of each FF and FcoR gene (X-axis) levels are shown. Each individual expresses a unique repertoire of FF and FcoR genes, and expression levels are stable following multiple cycles of ablation/regeneration.

**Extended Data Fig. 9.**
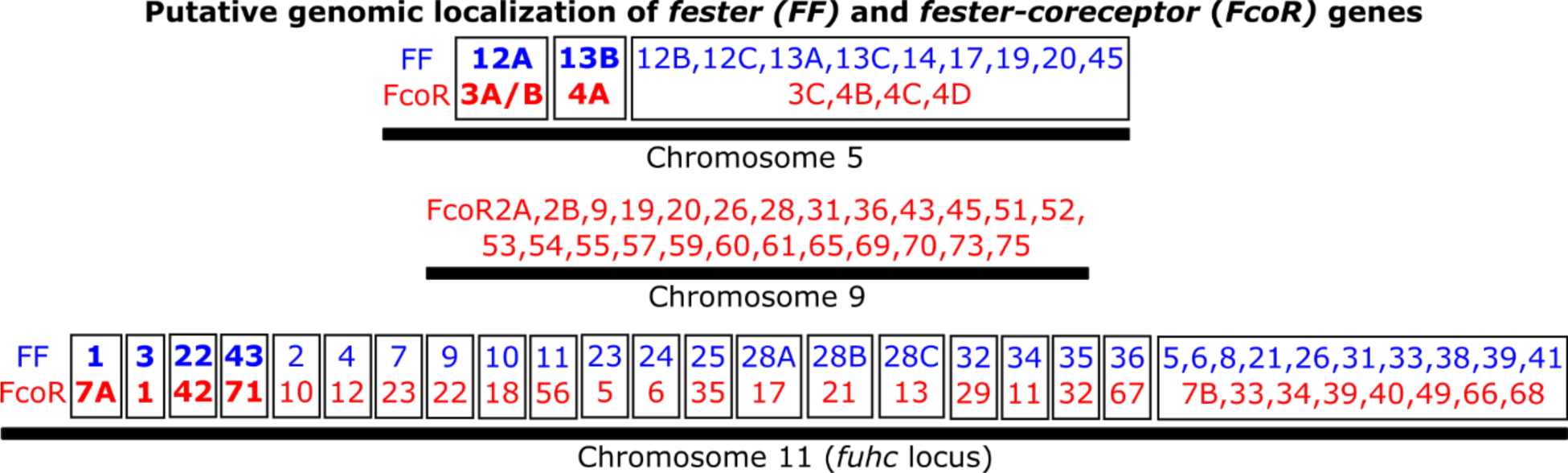
Predicted genomic localization of *fester* and *FcoR* genes. Predicted genomic localizations were estimated based on phylogenetic similarity between genes and chromosomal regions. Gene pairs were established using both phylogenetic, genomic and transcriptomic data. *fester* and *FcoR* genes are highlighted in blue and red, respectively. The genomic localizations of the genes highlighted in bold have been confirmed.

**Extended Data Fig. 10.**
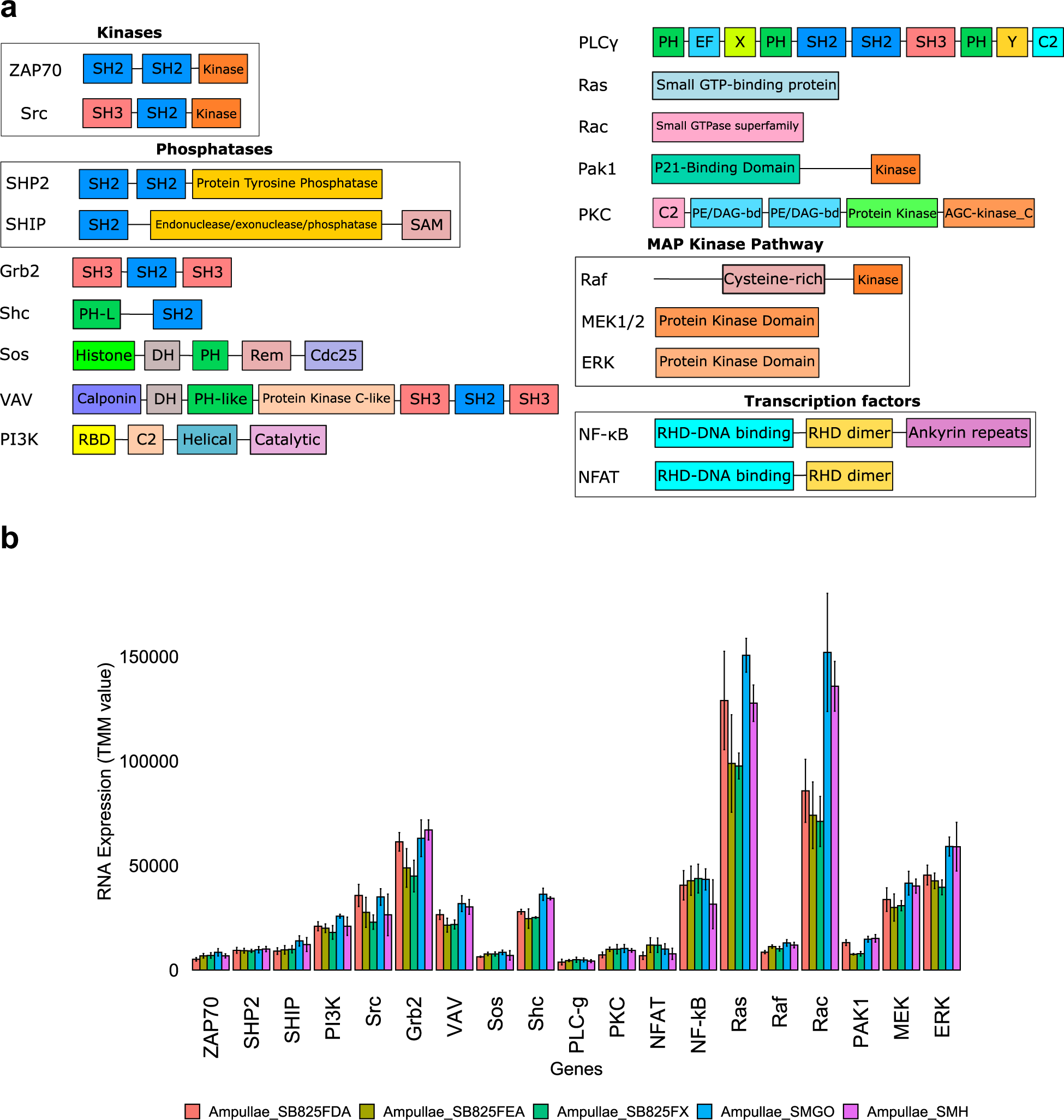
Structure and expression of proteins involved in ITAM and ITIM signaling from *B. schlosseri*. **a,** domain architecture of homologs of the indicated genes were identified in the transcriptome of *B. schlosseri*. **b,** expression of these genes was quantified in the vasculature system of five *B. schlosseri* individuals. Genes are plotted in the x-axis and RNA expression (TMM units) is plotted in the y-axis. Means and standard deviations were calculated between three or four replicates per individual.

**Extended Data Table 1.** Transcriptomes of *B. schlosseri* analyzed in this study.

| **Transcriptomes** | **Individuals** | **Accession number** |
| --- | --- | --- |
| Fertility-Blastogenesis | Fertile841, Fertile801, Infer001, Infer802, Mix Fertile, Mix Infertile | PRJNA263209 |
| Aging | SB825, SMH, SMGO | PRJNA474433 |
| Whole Body | SB825, SMH, SMGO, SKI | De Tomaso Lab (Jon Schultz). This study |
| Male | MDR, M001, MADM | De Tomaso Lab (Delany Rodriguez). This study |
| Vasculature (Ampullae) | SB825-FDA, SB825-FEA, SB825-FX, SMGO, SMH | De Tomaso Lab (Daryl Taketa). This study |
| ALDH Winner Loser | Loser 1, Loser 7, Winner 2B, Winner 6 | De Tomaso Lab. This study |
| BSA Vascular Cells (Ampullae) | Not specified | PMC3988187 |
| Blood | FL3, FLLC2, LFC4 | De Tomaso Lab (Shambhavi Singh). This study |
| IA6 Winner Loser | 81A, 81C | De Tomaso Lab. This study |
| Whole-body regeneration (WBR) | Isogenic strain | PRJNA798022 |

**Extended Data Table 2.**
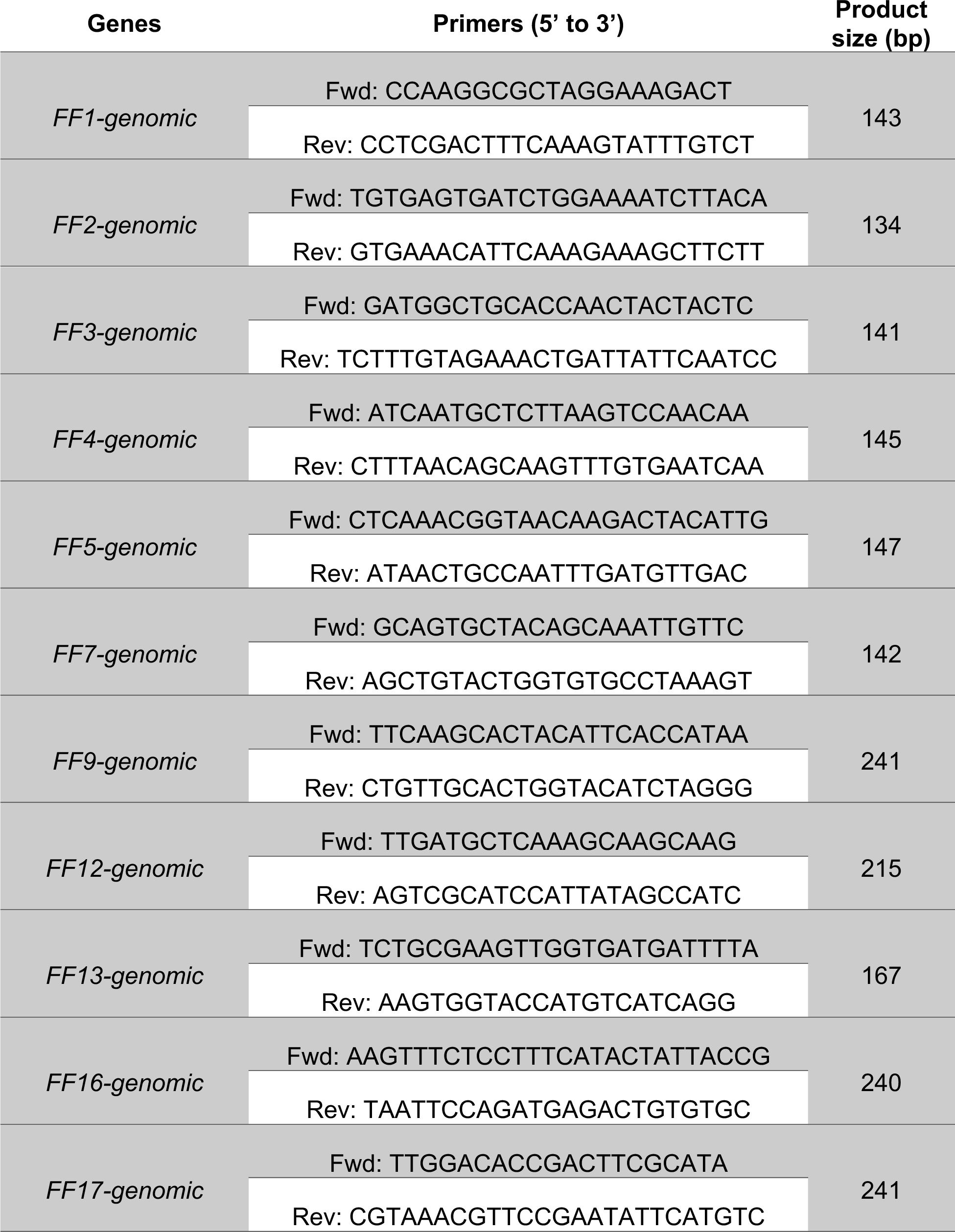
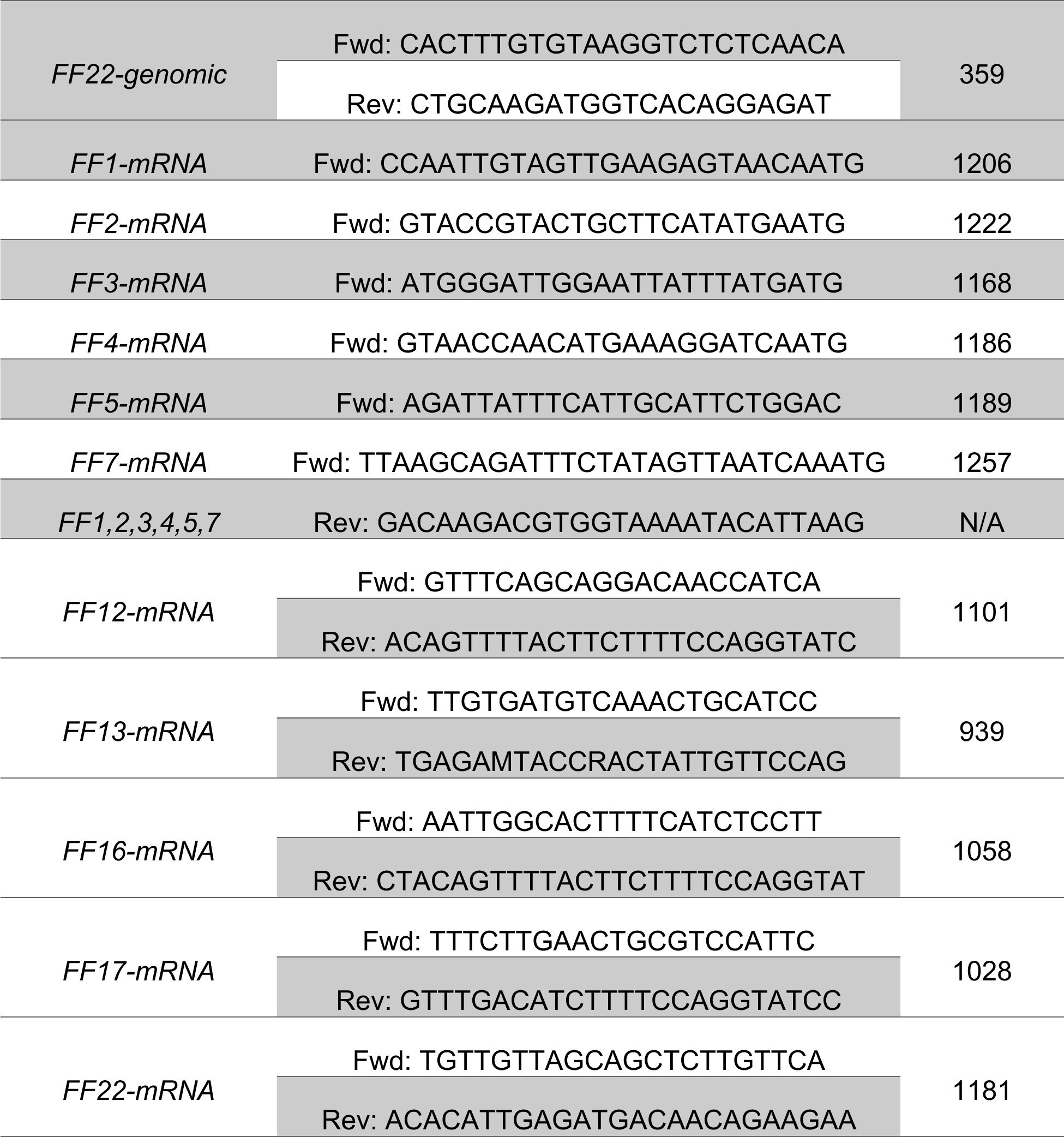
Primers used in this study for *B. schlosseri.*

**Extended Data Table 3.**
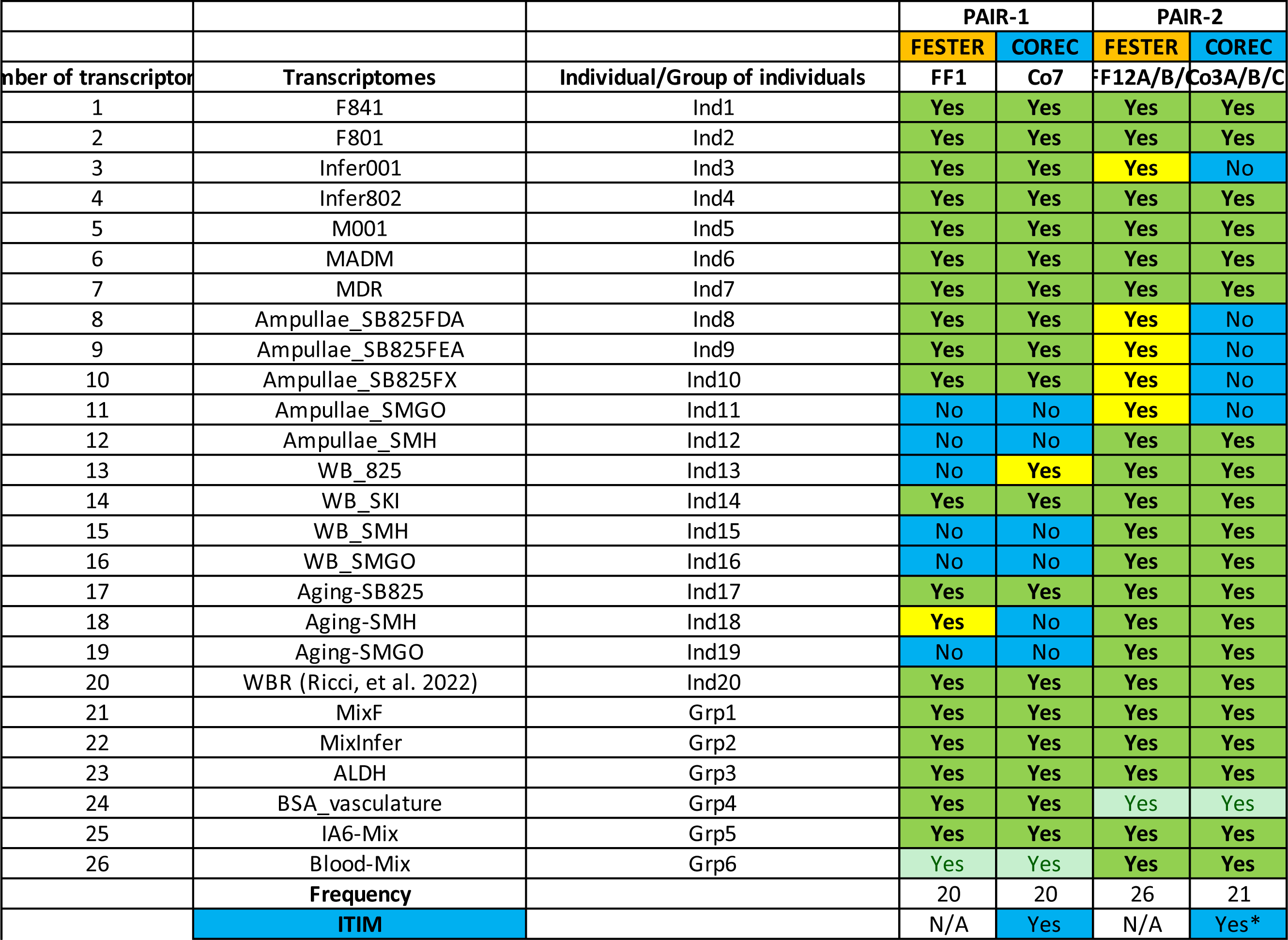
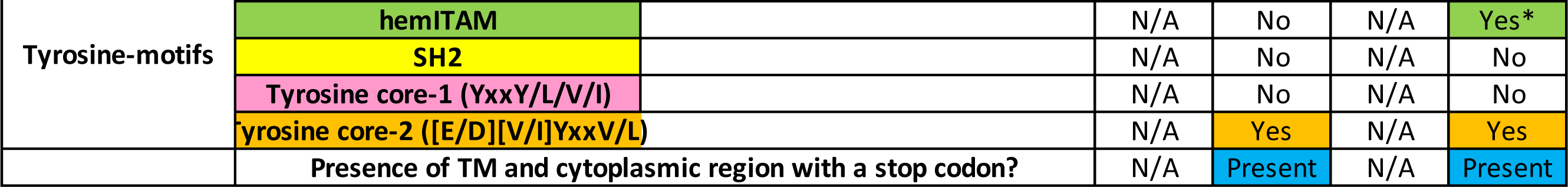
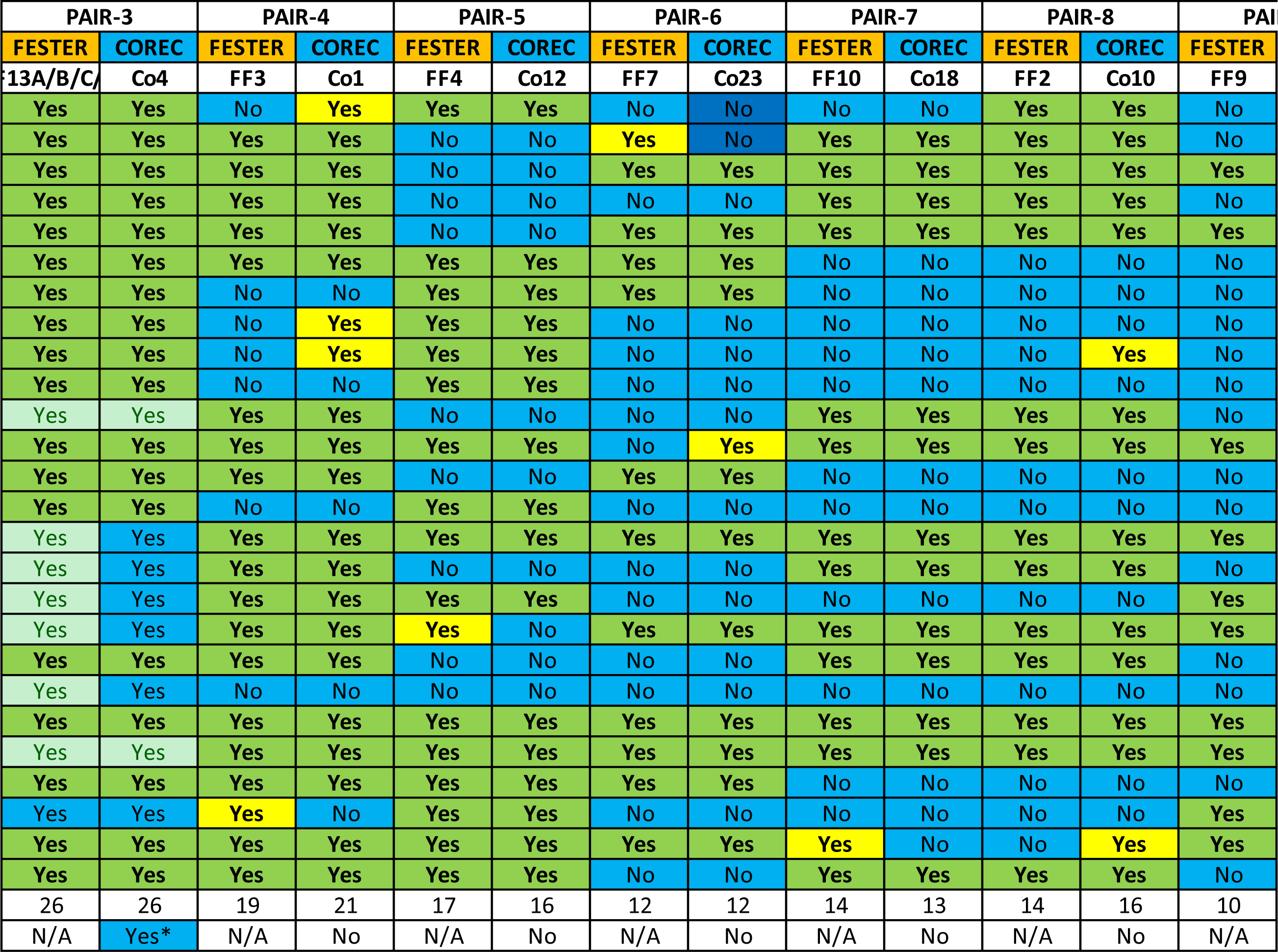
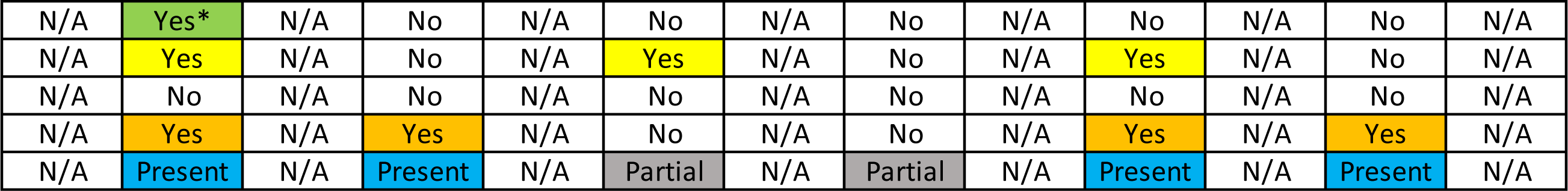
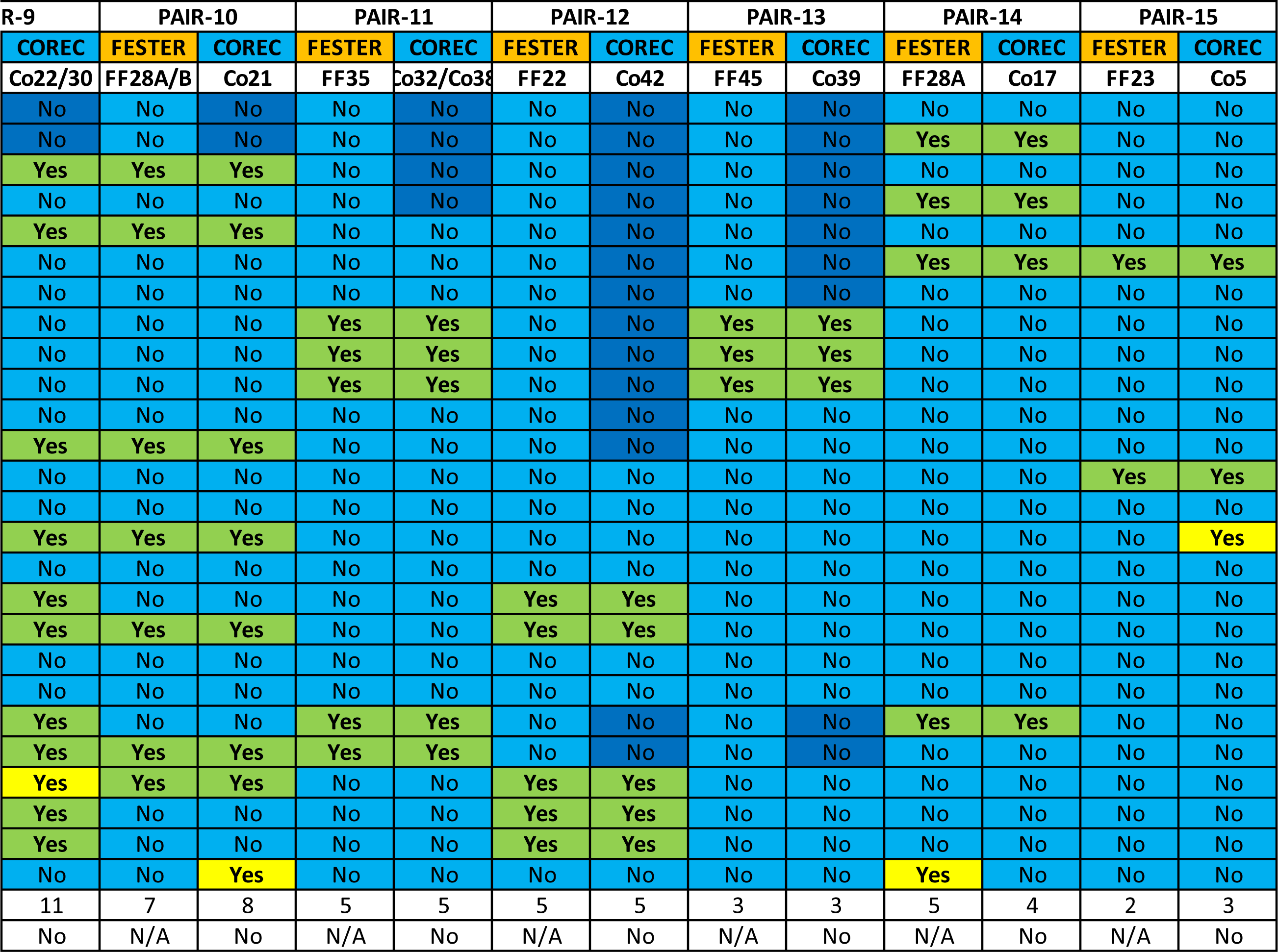
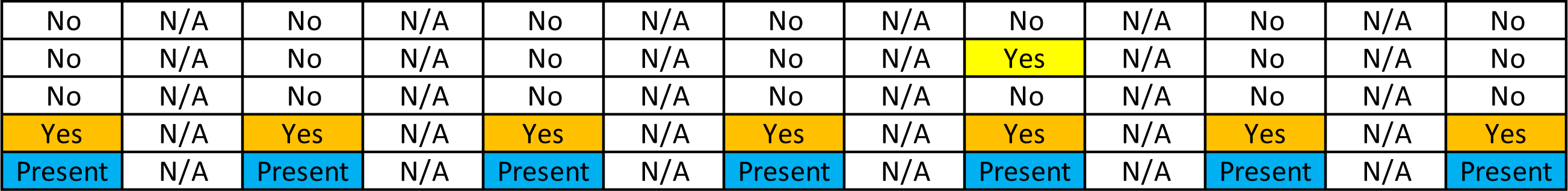
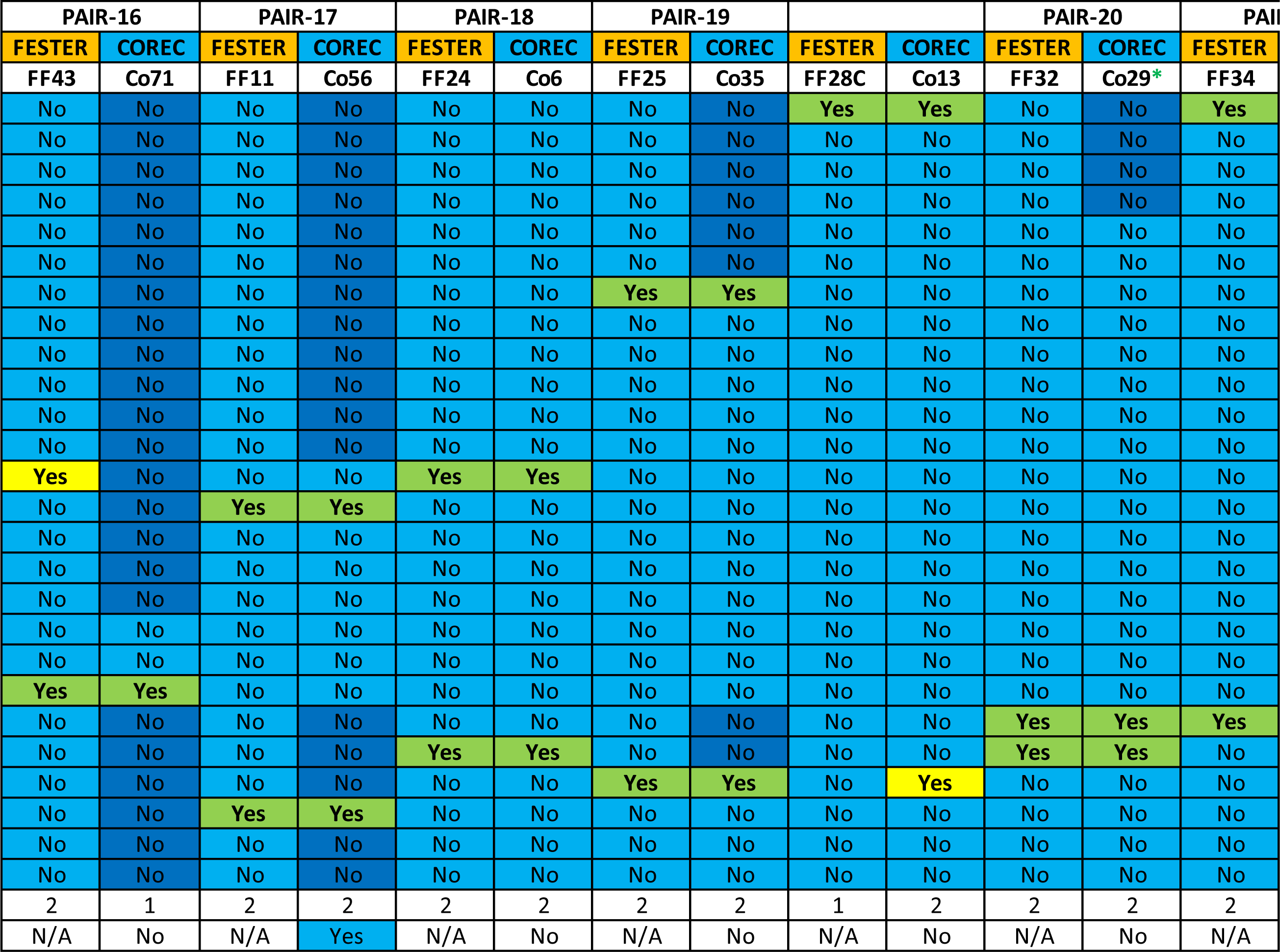
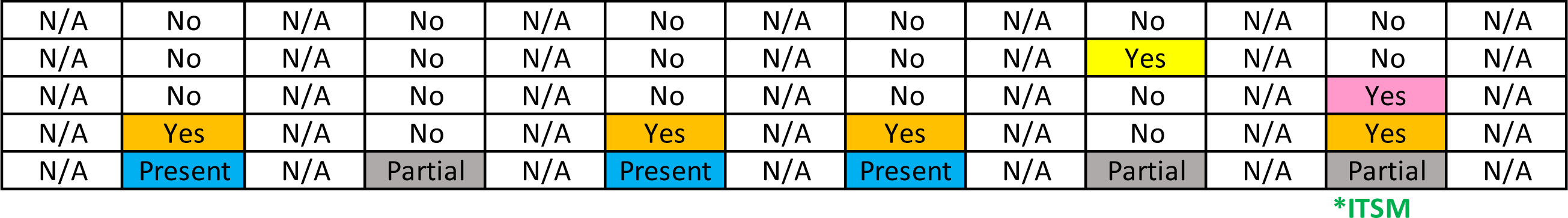
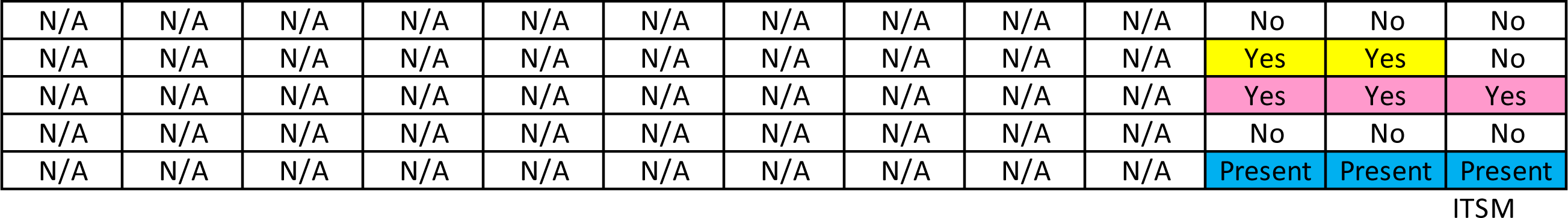
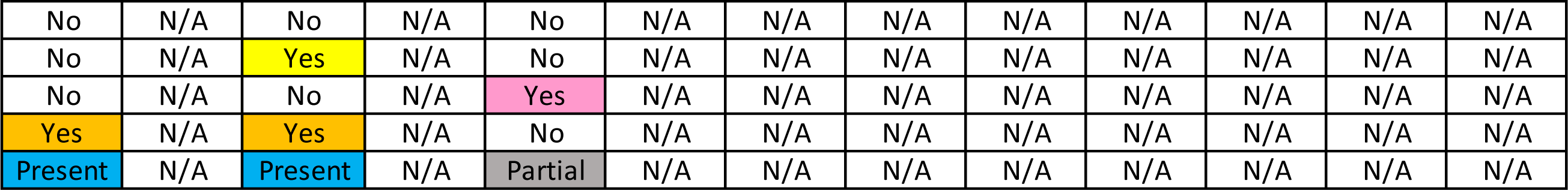
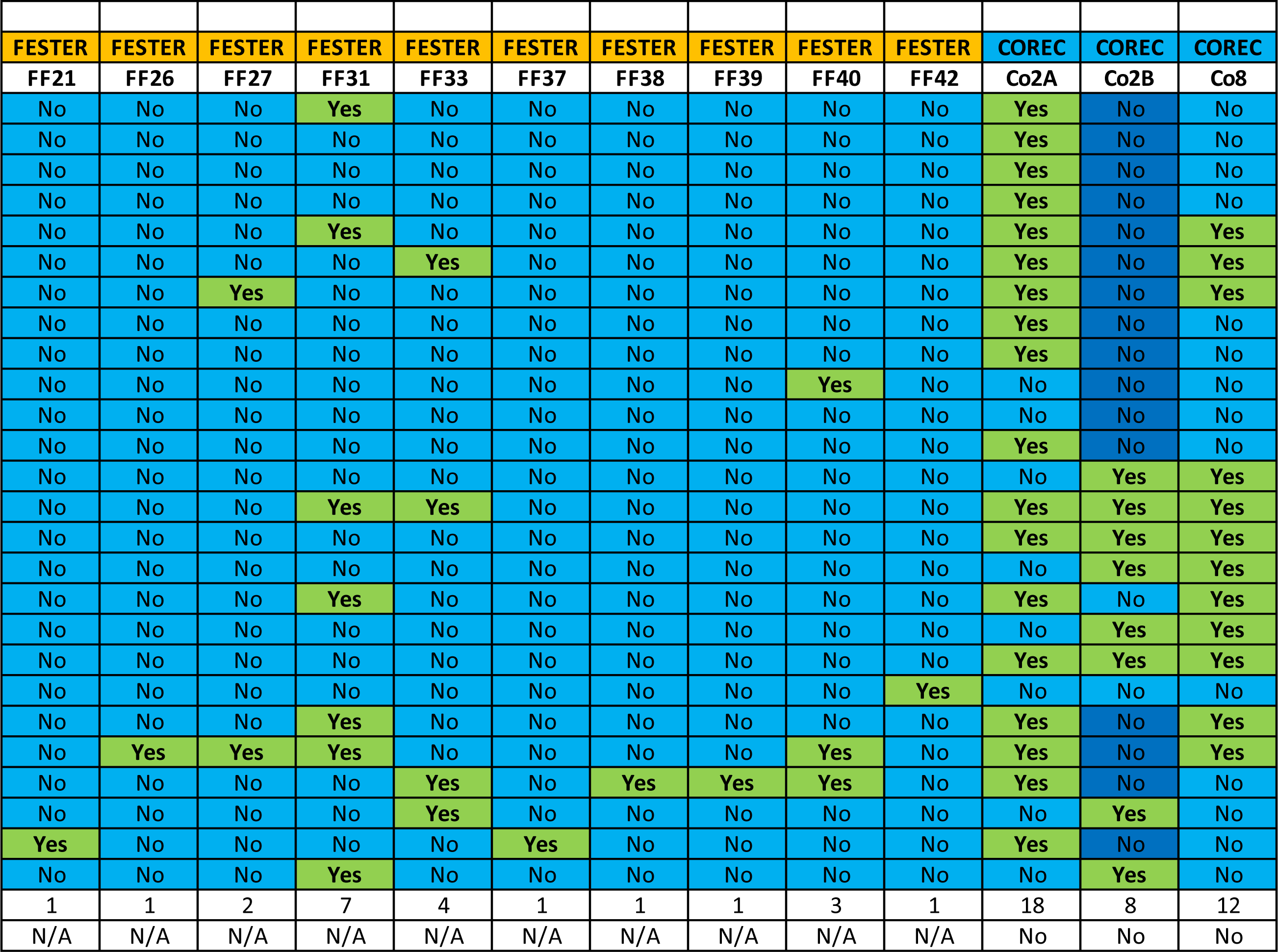
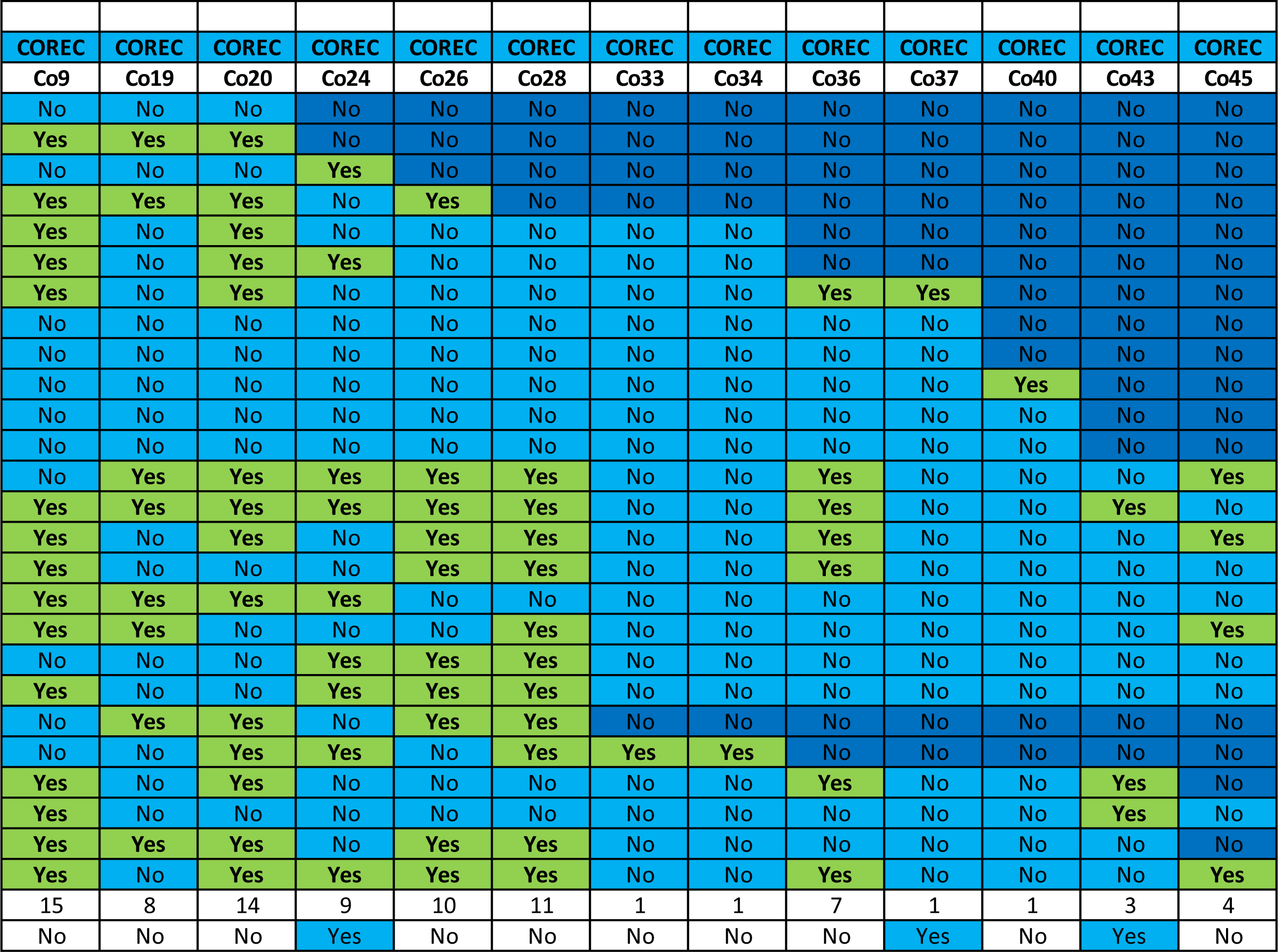
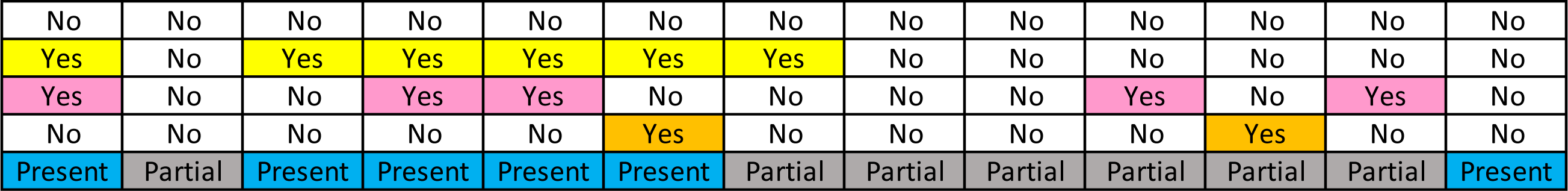
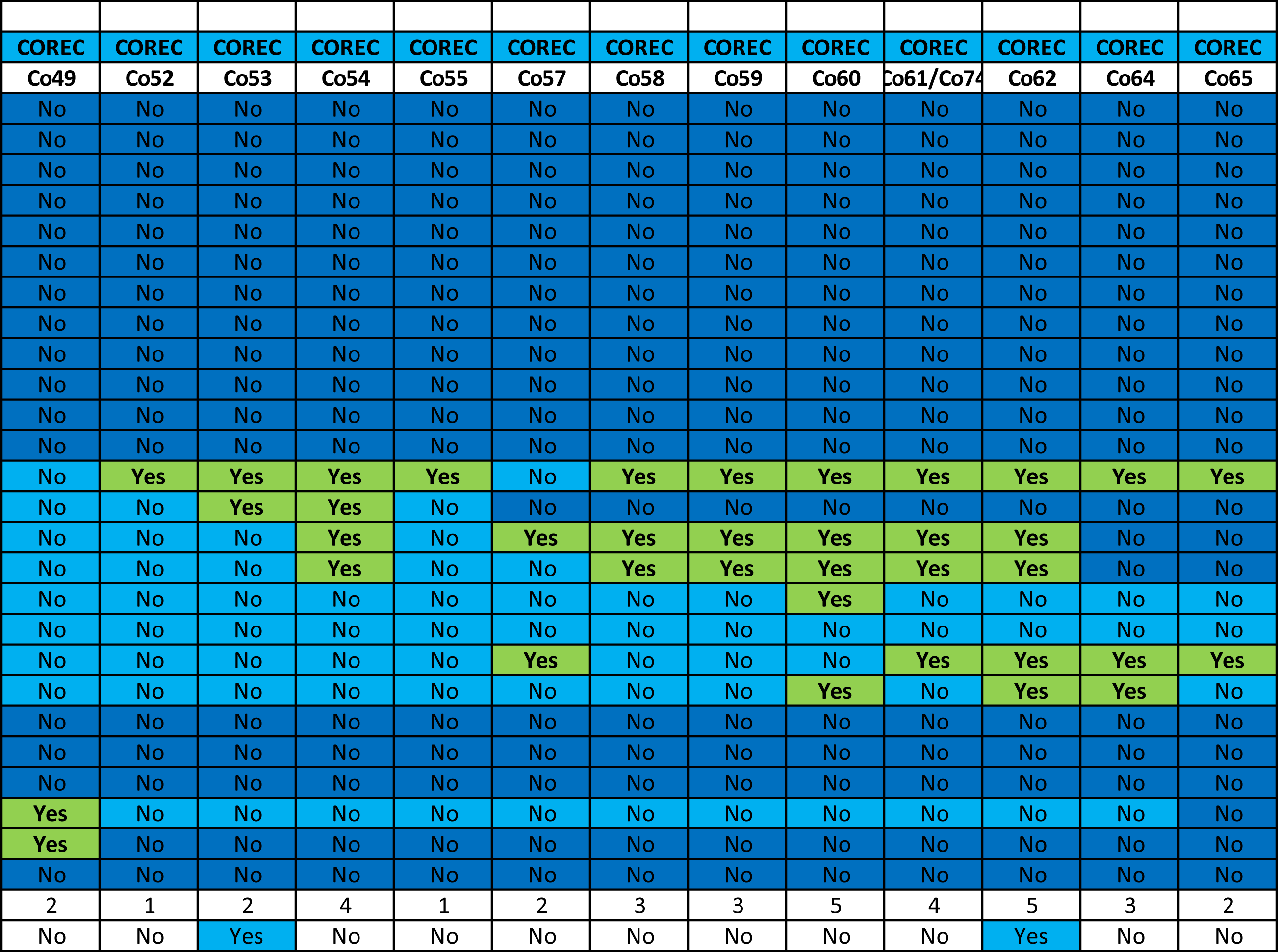
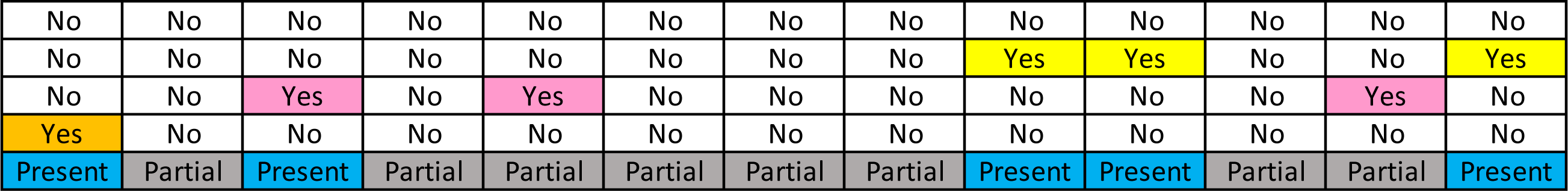
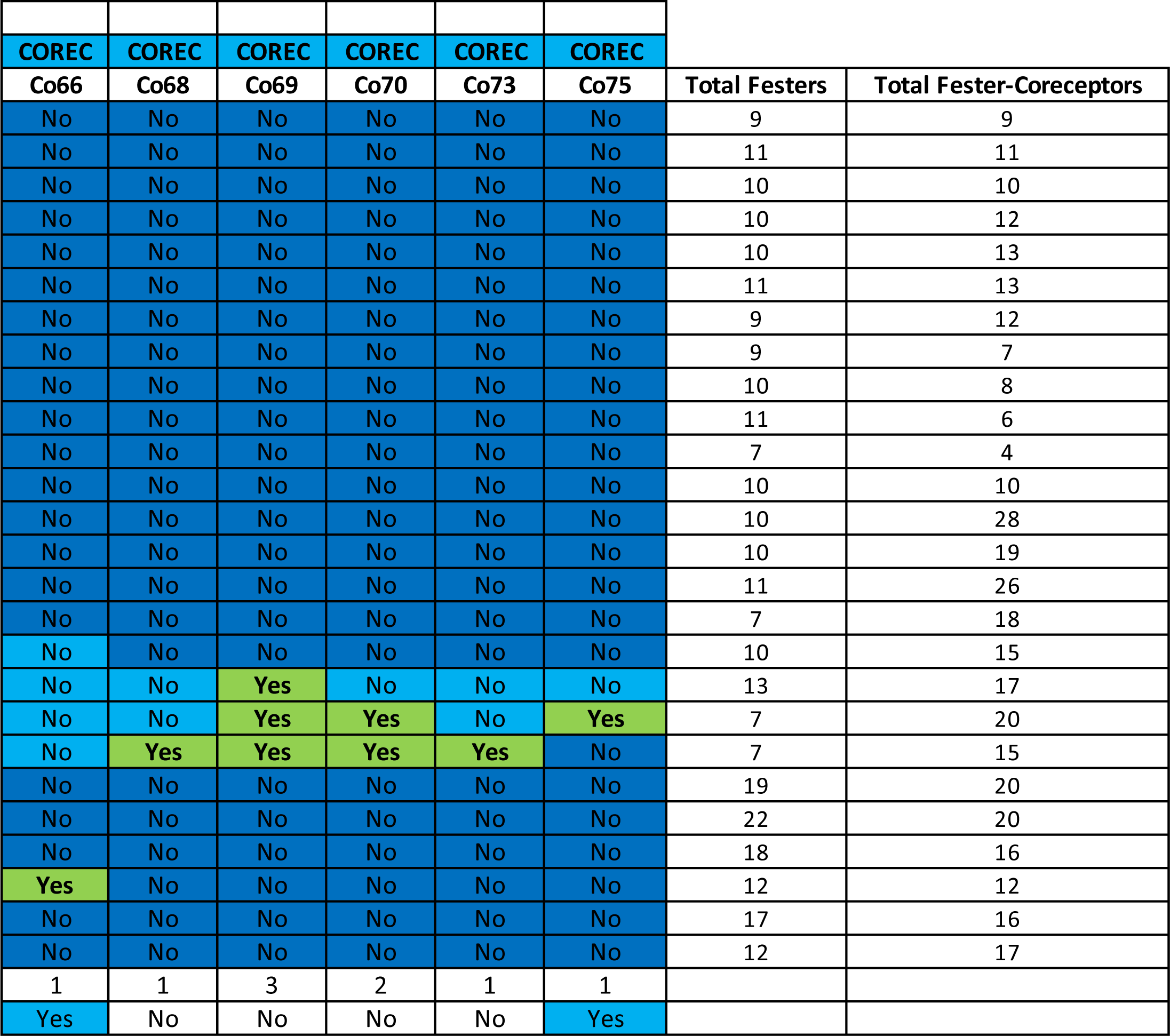
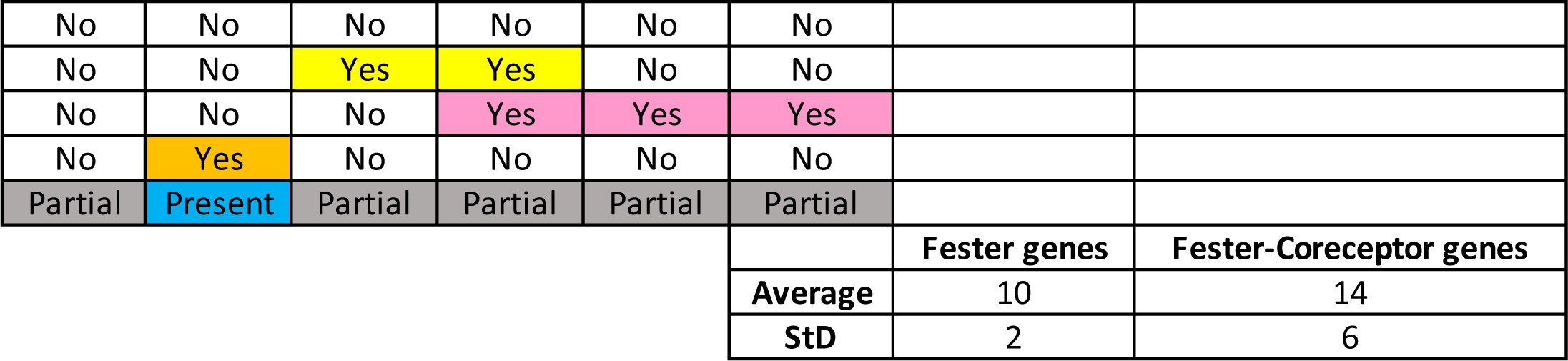
Is a converted Excel file over the next 18 pages.

**Extended Data Table 4.** Alternative splicing of *fester* genes.

| **Fester genes** | **Number of alternative splice exons between Sushi and TM-I Domains** | **Number of alternative splice exons in the cytoplasmic region** |
| --- | --- | --- |
| **FF1** | 2 | Not detected |
| **FF2** | 2 | Not detected |
| **FF3** | 2 | Not detected |
| **FF4** | 2 | Not detected |
| **FF5** | 1 | Not detected |
| **FF6** | 2 | Not detected |
| **FF7** | 3 | Not detected |
| **FF8** | 2 | Not detected |
| **FF9** | 2 | Not detected |
| **FF10** | 1-2 | 1 |
| **FF11** | 1 | 1 |
| **FF12/18/44** | 1 | Not detected |
| **FF13/15/16/19** | 1 | Not detected |
| **FF14** | 1 | Not detected |
| **FF17** | 1 | Not detected |
| **FF20** | Not detected | Not detected |
| **FF21** | 2 | Not detected |
| **FF22** | 2 | 2 (Introns) |
| **FF23** | 2 | 1 |
| **FF24** | 2 | 1 |
| **FF25** | Not detected | Not detected |
| **FF26** | 2 | 1 |
| **FF27** | Not detected | Not detected |
| **FF28/29/30** | 2/1/Not detected | 1 |
| **FF31** | Not detected | Not detected |
| **FF32** | 2 | 2 |
| **FF33** | Not detected | Not detected |
| **FF34** | Not detected | 1 |
| **FF35** | 2 | 1 |
| **FF36** | Not detected | Not detected |
| **FF37** | 2 | 1 |
| **FF38** | Not detected | Not detected |
| **FF39** | Not detected | Not detected |
| **FF40** | Not detected | Not detected |
| **FF41** | 2 | Not detected |
| **FF42** | Not detected | Not detected |
| **FF43** | Not detected | Not detected |
| **FF45** | 1 | 1 |

